# Activity Patterns Structure Food Web Interactions Through Time

**DOI:** 10.64898/2026.05.20.726571

**Authors:** Alexa M Scott, Emily K Studd, Carling Bieg, Brett M Studden, Kevin S McCann, Bailey C McMeans

## Abstract

Many mobile animals move to locate and consume resources, making energy gain and growth dependent on activity. Yet the role of activity in shaping predator-prey interactions in food webs has not been broadly considered. Here, we synthesize empirical examples to examine how three activity traits (mean, variance, and timing) vary among taxa (fish, mammals, birds) and between predators and prey across temporal scales. We then use predator-prey models to explore how these diverse ‘activity patterns’ influence stability. Motivated by emerging activity patterns, our theory shows that fluctuating activity rates can drive predator-prey interaction strengths with major consequences for stability. Future research is needed on activity trait patterning, links between activity and attack rates, and the consequences of activity for predator-prey interactions to whole food webs. This is especially critical as human-driven changes to abiotic cues increasingly alter animal activity rates and may rewire food webs.

## Introduction

The majority of earth’s ecosystems experience cycles in abiotic conditions at multiple scales from hours to days to months to years (figure 1). The 24-hour daily cycle of light and dark is highly predictable and a dominant abiotic cue for biological rhythms and diel activity patterns (figure 1a) (Bradshaw and Holzapfel 2007). Fluctuations in moonlight are similarly predictable every 30 days, illuminating the landscape during the full moon phase and immersing the landscape into darkness during the new moon phase (figure 1b). At the seasonal scale, cyclical variation in environmental conditions are created by the earth’s rotation around the sun, breaking the year up into distinct seasons (figure 1c). Across these timescales, different abiotic signals can fluctuate through time together (synchronously), like temperature and solar irradiance (figure 1a), or in opposition with each other (asynchronously), like temperature and snow depth (figure 1c), creating a heterogeneous and diverse poly-rhythmic environment experienced by organisms through time (Zhu et al. 2019, Bernhardt et al. 2020, von der Heydt et al. 2021).

**Figure 1:**
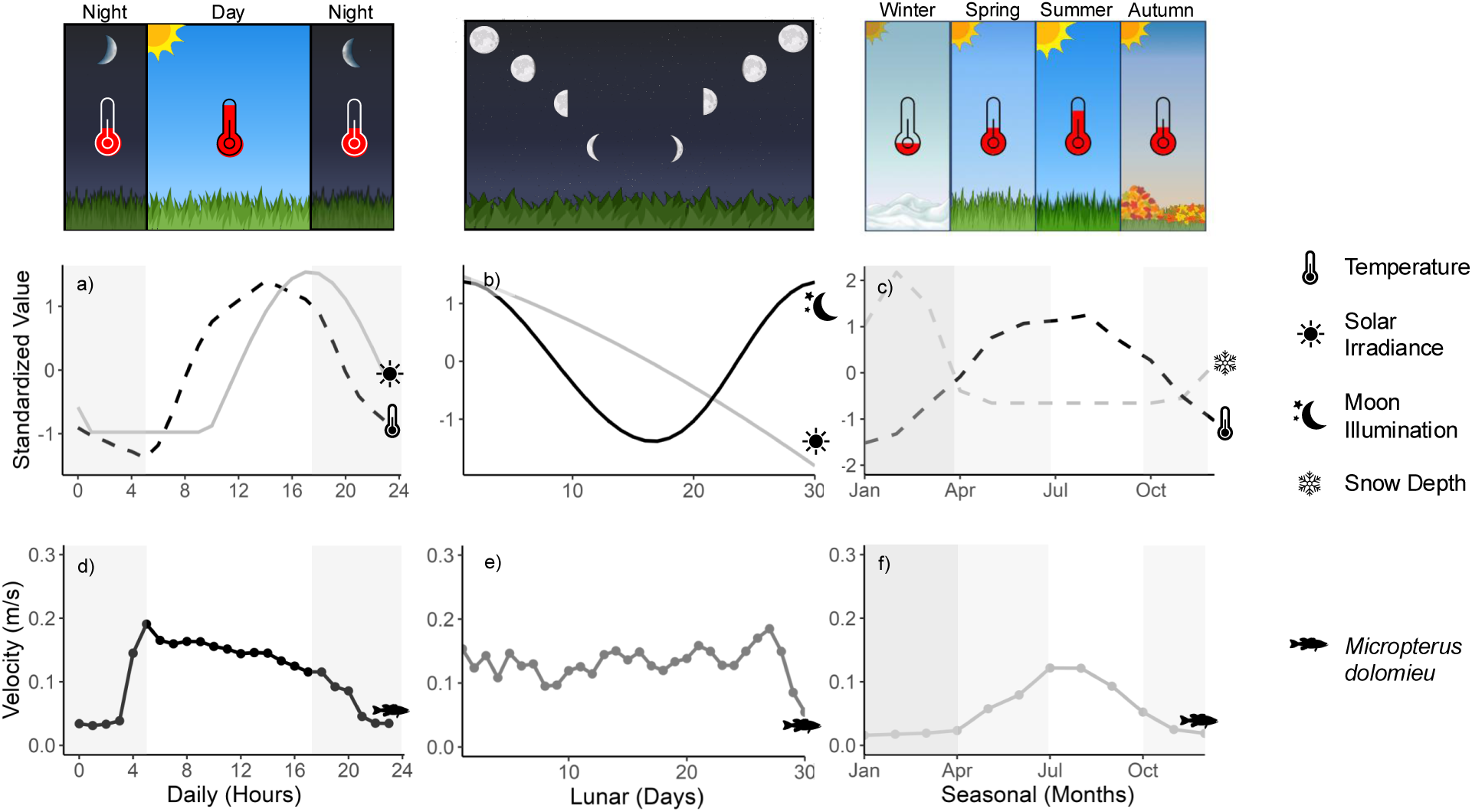
The fluctuation of various abiotic signals across the a) daily (hours), b) lunar (days), and c) seasonal (months) timescales, and the activity patterns of the smallmouth bass (*Micropterus dolomieu*) across the d) daily, e) lunar, and f) seasonal timescales. Original sources for all data can be found in table S2.

Temporal fluctuations in the environment incite a suite of physiological and behavioural responses in organisms (Kronfeld-Schor et al. 2017, Bernhardt et al. 2020). Activity, which we define here as the movement rate (e.g., distance/time) of a whole organism through space, is a widespread response to environmental periodicities across diverse taxa. In mobile animals, activity is often required to locate and consume resources (Elliott et al. 2009), to reproduce (Schmidt et al. 2009), disperse(Arndt and Evans 2022), or thermoregulate (Maloney et al. 2005). On the other hand, inactivity can arise to save energy (Lesku et al. 2012), decouple from harsh environments (Chesson and Huntly 1997), to sleep (Rattenborg et al. 2017), and to avoid predators (Lima and Dill 1990, Tambling et al. 2015) or competitors (Monterroso et al. 2016). The timing of activity is diverse in nature as species are differentially sensitive to abiotic cues, thus targeting their activity towards different times of the day, month, and year (for example, see activity of *Micropterus dolomieu* in figure 1d, 1e, 1f). Notably, the diverse timing of activity allows organisms to partition time as an ecological resource (Kronfeld-Schor and Dayan 2003, Kronfeld-Schor et al. 2017) and can promote coexistence among competitors in a community (Chesson 2000, Hood et al. 2021). In food webs, prey are known to time their activity to minimize encounters with predators (Lima and Bednekoff 1999, Tambling et al. 2015), while predators can synchronize their activity with prey (Lang et al. 2019) or with abiotic signals (Bosiger and McCormick 2014), suggesting a delicate balance between optimal activity strategies for interacting species. However, the role of activity in predators and prey – and specifically, how the covariance between activity patterns shapes predator-prey interactions – has not been as widely studied in food web ecology as other mechanisms like density or spatial overlap (Lima 2002).

Recent advances in tracking technologies allow for relatively continuous measurement of activity patterns at the daily, monthly, seasonal, and multi-annual scales (Hussey et al. 2015, Kays et al. 2015, Hofman et al. 2019). Movement ecology is its own subfield of ecology that considers the how, why, when, and where of an organisms’ movement (Nathan 2008). Given the abundance of movement behavior data now being compiled with the increased use of GPS and telemetry, and the multitude of emerging activity patterns (figure 2), there is significant potential to advance our understanding of time varying activity rates, attack rates and interaction strengths (flux based; Nilsson and McCann 2016). The general consequences of this time-varying predator and prey activity rates for predator-prey attack rates, food web structure, and dynamics is an important but underexplored research question.

**Figure 2:**
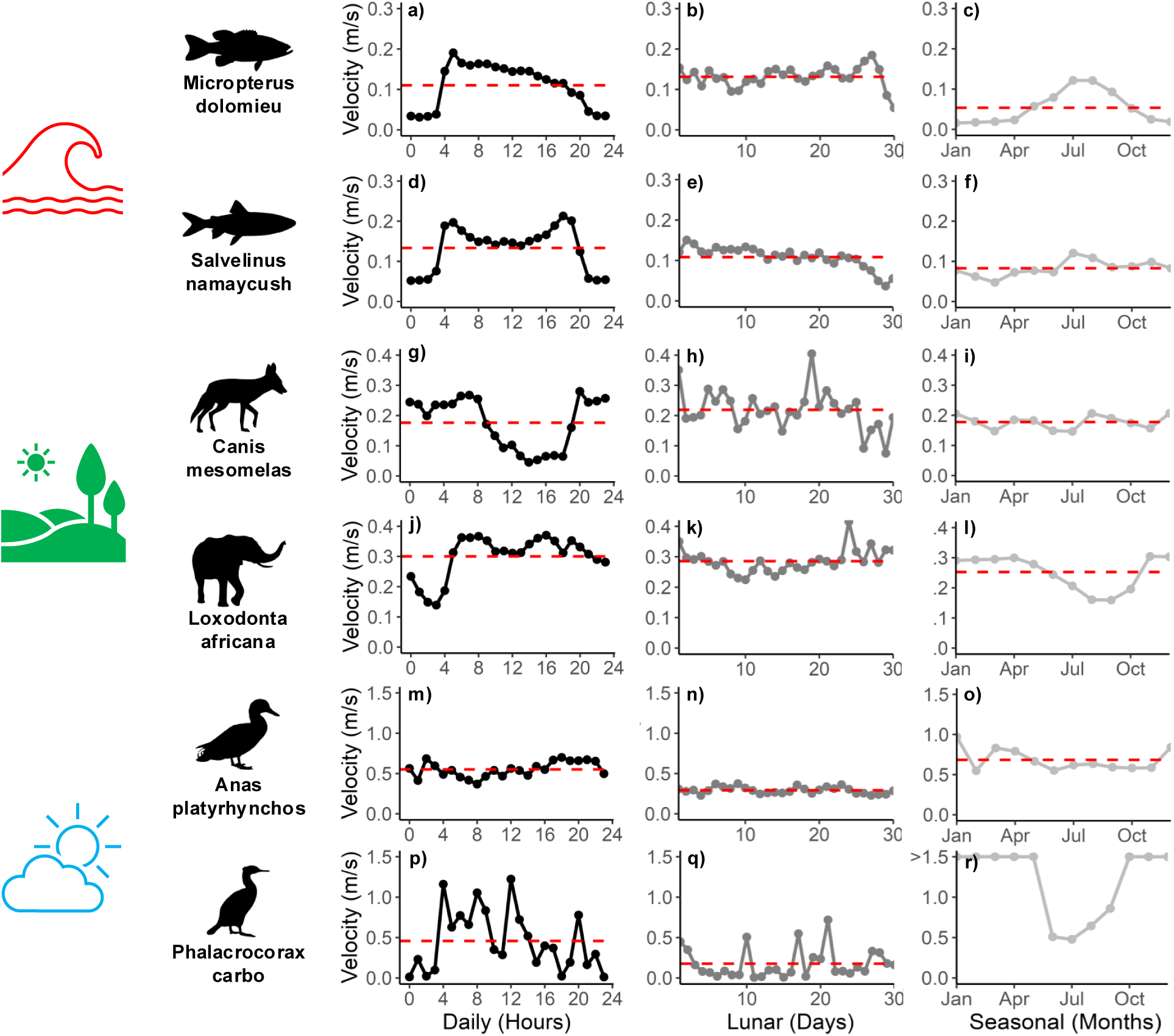
Activity patterns of various species across the daily (black lines), lunar (grey lines), and seasonal (light-grey lines) timescales with their corresponding activity mean (red dashed line). Small mouth bass (*Micropterua dolomieu*; a-c) and lake trout (*Salvelinus namaycush*: d-e) are both aquatic cohabitants from Algonquin Park, Ontario Canada. Both terrestrial species, the black-backed jackal (*Canis mesomelas*: g-i) and the African elephant (*Loxodonta africana*: j-l), reside in Etosha National Park, Namibia, Africa, and the mallard duck (*Anas platyrhynchos*) and the great cormorant (*Phalacrocorax carbo*) are both aerial birds found near Lake Constance, Baden-Württemberg, Germany. Note, the great cormorant’s seasonal mean (3.237 m/s), exceeds the axis range set to highlight activity variability within each timescale due to the species migration periods spanning from October to April (see figure S3.1 for full profile). Original sources for all data can be found in table S2, and each species’ tagging duration can be found in table S3.

Towards this, we argue that activity patterns in predators and prey should play an important role in shaping attack rates and interaction strengths within food webs. Using empirical examples, we begin by highlighting the diversity in activity patterns at multiple temporal scales and how predators and prey vary in their activity patterns. We then theoretically explore the consequences of different activity patterns in predators and prey for predator-prey dynamics, before raising new directions and questions for future research. Our goal is to motivate future empirical and theoretical work and to make the case that activity should be viewed as a fundamental aspect influencing nature’s trophic relationships and therefore the functioning and resilience of whole food webs. Effectively, we ask ecologists to consider not only ‘where’ but ‘when’ organisms are active or inactive during a day, month, and year, and how this affects their interactions with other organisms in a food web. In what follows, we focus on three attributes of activity – **mean, variability** and **timing** - and three timescales – **daily (period = 24 hours), lunar (period = 30 days),** and **seasonal (period = 12 months)** – as representations of key rhythms for biotic organization (figure 1). Additionally, we emphasize that by considering the nature of activity patterning broadly across scales, this approach provides the scaffolding for an effective integration of movement and food web ecology. Finally, given that the timing of many abiotic signals are shifting under climate change at daily (Gilbert et al. 2023) and longer timescales (Wolkovich et al. 2014), there is a pressing need to consider the general consequences of activity patterns for the structure and stability of food webs.

### Diverse activity patterns across taxa and timescales

To look at the potential diversity of activity patterns across taxa and timescales, we gathered data from six species (three taxonomic and cohabitant pairs: see methods in supplementary material) that had empirical activity data at the daily, lunar, and seasonal timescales and quantified the mean, variance, and timing of activity over each timescale. For each taxonomic and cohabitant pair, the month in which both species displayed the highest overall daily activity was selected for daily-scale comparison, while the month with the highest average daily activity during the lunar cycle was chosen for lunar-scale comparison (see supplementary material S1.3). We see this as a first step towards the documentation of animal activity strategies, and their implications, in a manner akin to how ecologists have categorized organismal life history strategies.

### Activity mean and variance

The organisms display a wide range of mean activity rates across species and taxa at all timescales. For instance, at the daily scale, movement rates increase from fish to mammals and finally to birds who have approximately five times the movement rate of the aquatic species (Fig. 3a) – this pattern is largely maintained across all temporal scales. To a lesser extent, variation exists within taxonomic groups. African elephants (*Loxodonta africana*), for example, have 1.2 to 1.5 times the mean movement rate of the black-backed jackal (*Canis mesomelas*) across all temporal scales (Fig. S3.2a) while smallmouth bass (*Micropterua dolomieu*) and lake trout (*Salvelinus namaycush*) have similar mean activity rates, with which species is more active being dependent on the temporal scale (Fig. 3d). By comparing figure 2c and 2f, these figures suggest that coldwater-adapted lake trout move at greater rates in the winter and lower rates in the summer relative to warmwater-adapted smallmouth bass. Finally, the mallard duck (*Anas platyrhynchos*) can have a faster mean activity rate across shorter timescales (daily, lunar), and a slower mean activity rate across longer timescales compared to the great cormorant (*Phalacrocorax carbo*) (Fig S3.3a).

**Figure 3:**
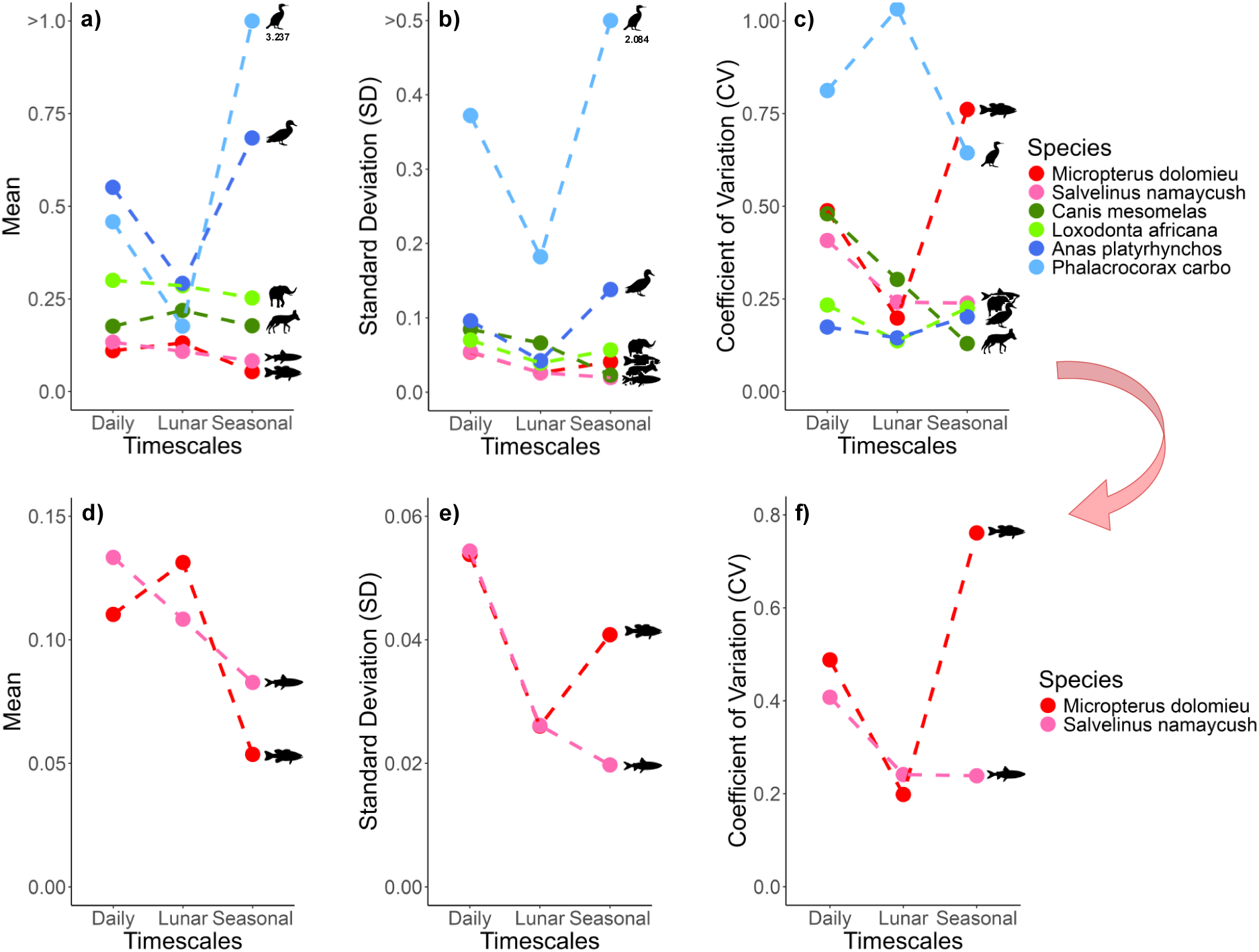
Relatively faster and slower activity patterns (measured by (a) mean) and variability of the activity patterns (measured by (b) standard deviation (SD) and (c) coefficient of variation (CV)) of various species across the daily, lunar, and seasonal timescales. Close up on the two aquatic species differing in their activity d) mean, e) standard deviation, and f) coefficient of variation across each timescale. Close ups on the terrestrial and aerial taxonomic pairs can be found in figure S3.2 and figure S3.3 respectively. Original sources for all data can be found in table S2.

A wide range of variance in activity is also apparent among species, measured as both standard deviation (SD, figure 3b) and coefficient of variation (CV; calculated as SD relative to a species’ mean, figure 3c), again highlighting the diversity of potential temporal activity patterns. We note here that while the mean, SD, and CV are comparable across all species (Fig 3a-c respectively), the element in which each species moves in (i.e., water, land, air), may account for the appearance of a taxonomic pair having a much lower activity mean and standard deviation (i.e., both aquatic species, smallmouth bass and lake trout). However, when the mean, SD, and CV are examined within each taxonomic pair (for example, see aquatic cohabitants in figure 3d, 3e, 3f respectively), vast differences can be seen across all timescales (see figure S3.2 for terrestrial and figure S3.3 for aerial).

The great cormorant, a species with high mean activity (figure 3a, S3.3a), has relatively high activity variance when measured as SD (figure 3b, S3.3b), but lower activity variance when measured as CV (i.e., once this large SD is expressed relative to the high mean, figure 3c, S3.3c). Alternatively, species like smallmouth bass, who have a lower mean activity (figure 3a, 3d) and a lower variance when measured as SD (figure 3b, 3e), have a relatively high activity variation when expressed as CV (figure 3c, 3f). The lower CV of the mallard duck reflects their relatively constant activity throughout the day, month and year (figure 2m, 2n, 2o respectively), while the higher CV of smallmouth bass reflects periods of increased and decreased activity arising at the daily, lunar and seasonal scale (figure 2a, 2b, 2c respectively).

While there are considerable differences among species’ activity mean and variation within each timescale, there is also notable variability across timescales. For example, black-backed jackal have relatively high activity variance at the daily and lunar scales (both SD and CV, figure 3b, S3.2b and 3c, S3.2c respectively) but reduced activity variance at the seasonal scale (figure 3b,c, S3.2b,c) due to constant activity all year round (figure 2i).

For most species, other than the black-backed jackal and the great cormorant, the standard deviation in activity relative to the mean activity (i.e., CV), tended to have lower variation at the lunar scale relative to the daily and seasonal timescales (figure 3c). This result may suggest that abiotic periodicities at the monthly scale do not drive activity patterns as dramatically or as generally as the night-day cycles or the seasonal cycle (e.g., summer-winter, wet-dry), however, more work is needed in a broader range of taxa to explore this idea as potential exceptions could exist (i.e., marine species experiencing tides; Palmer 1973).

### Timing of activity

While activity mean and variability tell us how fast vs. slow and constant vs. variable an organism’s movement is, activity timing tells us *when* an organism is active at the daily, monthly and seasonal scale (figure 4a, 4b, 4c). This additional layer is necessary for understanding how temporal activity patterns are structured and whether activity is synchronized or asynchronized among potentially interacting species. Synchronized activity, for example, may enhance attack rates as both species are active simultaneously. At the daily scale, there are four common strategies in which species time their activity: diurnal (active during the day), nocturnal (active during the night), crepuscular (active at dawn and dusk), and cathemeral (irregular activity throughout the day and night) (Tattersall 1987, Maor et al. 2017). From our examples in figure 2, diurnal species include the smallmouth bass, the lake trout, and the great cormorant (figure 2a, 2g, 2p respectively), nocturnal species include the black-backed jackal (figure 2d), and cathemeral species include the mallard duck and the African elephant (figure 2m, 2j respectively). Notably, these different activity patterns mean that, among different species, activity can be synchronous or asynchronous. In figure 4, we classify the timing of each species’ activity across timescales relative to the mean movement rate, using a t-based interval around the mean (± t·SE) to distinguish lower (tan), average (gray), and higher (black) activity levels. Here, species are synchronized when maximum activity times (i.e., black areas) are aligned, and asynchronized when activity is timed differently (i.e., black and tan areas align). At the daily scale, figure 4a suggests that the two fish species, *Micropterus dolomieu* and *Salbelinus namaycush*, are largely aligned (synchronous) in time, whereas the terrestrial species, *Canis mesomelas* and *Loxodonta Africana*, diverge in their timing (asynchronous).

**Figure 4:**
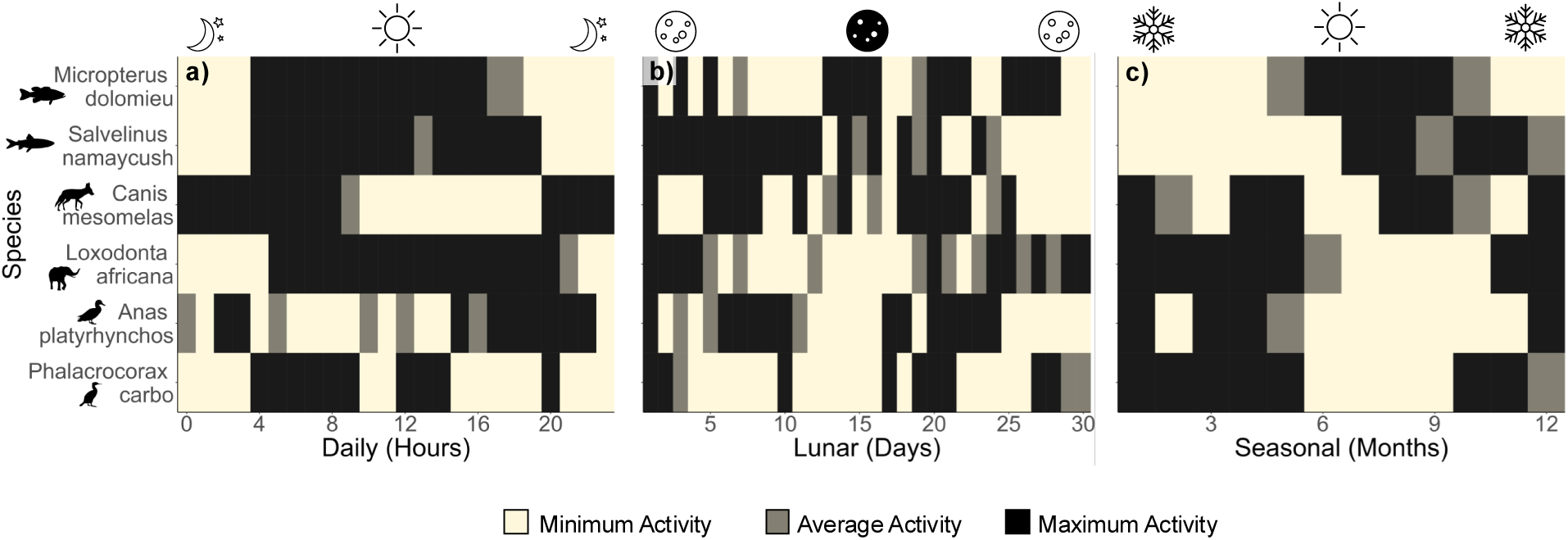
Minimum to maximum activity rates of species across the (a) daily, (b) lunar, and (c) seasonal timescales. Activity is classified relative to the mean movement rate using a t-based threshold (mean ± *t*·SE, where t = t_0.75,*df*_). Values below the lower bound (tan), within the interval (gray), and above the upper bound (black) represent minimum, average, and maximum activity, respectively. Species show both synchrony and asynchrony of activity rates at all timescales. Original sources for all data can be found in table S2.

At the lunar scale, changes in illumination from moonlight may encourage species to alter the timing of their foraging behaviours during different phases of the moon (figure 4b). On bright nights, nocturnal species may increase their foraging activity as their prey become easier to locate (Kronfeld-Schor et al. 2013). However, prey may reduce their own activity to reduce predation risk during the full moon phase (Kronfeld-Schor et al. 2013). The black-backed jackal, for instance, increase their foraging activity during each half-moon phase (first and last quarter), due to impaired vision when nights are completely dark, and increased detectability by prey during full moon phases (figure 2h, 4b; Ferguson et al. 1988). These diverse responses to moonlight set the stage for either synchrony or asynchrony to arise in activity timing at the lunar scale (figure 4b).

Finally, the predictable nature of seasonal cycles set the stage for species phenology, i.e., the timing of biological processes, including activity (Helm et al. 2013). Many organisms tend to experience maximum activity during the productive breeding periods (e.g., summer, wet) and minimum activity during the non-productive periods (e.g., winter, dry) at seasonal scales (figure 4c), although some remain active year-round in both aquatic and terrestrial ecosystems (McMeans et al. 2015, Humphries et al. 2017). Again, we see activity occurring at similar or different times of the seasonal cycle which synchronizes or asynchronizes activity patterns among species (figure 4c).

From the vast array of different activity strategies, the three key traits discussed here - mean activity rate, activity variance, and the timing of activity - likely have profound implications on species interactions, such as predator-prey interactions. In a sense, a species’ average activity sets the baseline around which variation occurs and represents the culmination of behaviour, metabolic strategies, life history and biotic interactions that ultimately mediate food web interactions in space and time. Animals that behaviourally respond to environmental cues – with predictable patterns of activity and inactivity timed around a day, month, or year – display considerable variation in their movement rates within these timescales, while others take on a more consistent pattern of activity over time, relative to their baseline (i.e., average) movement (figure 3b, 3c). While the timing of different behavioural patterns has been documented, this has generally been done with timescales being considered in isolation (as opposed to a more general framework for classifying activity patterns across scales, as we suggest here), and not done in a way that clearly links to the structure and function of communities/food webs. Importantly, all these different patterns of organismal responses to varying conditions at different scales suggest that activity patterns (and therefore predator-prey interactions) may vary in ways that could either synchronize activity rates in time (i.e., predator and prey simultaneously active) or decouple predator-prey interactions (e.g., one species inactive while another is active). Therefore, understanding temporal patterns in species’ activity and covariance between interacting species is a necessary first step towards understanding how species interactions (i.e., encounters in space) are structured through time.

### Predator-prey activity patterns

Differences in the mean, variation, and timing of activity between predators and prey could play a significant but poorly understood role in interaction strength and energy flow within food webs. Given the assumption that fast predator and prey activity rates imply stronger predator-prey attack rates (Öhlund et al. 2014), then higher mean activity rates in ‘faster’ species would be expected to increase prey encounters relative to predators that move more slowly or are more sedentary on average (Vander Vennen et al. 2016). Similarly, temporal variation in activity ought to significantly alter the overall strength of species interactions over time (e.g., time-averaged or - integrated consumption rates). For example, predators that are more constantly active, including cathemeral predators at the daily scale (Cox and Gaston 2024) and winter/dry active predators at the seasonal scale (McMeans et al. 2015, Humphries et al. 2017), should have more sustained interactions with prey compared to predators with more variable activity. Finally, given that predators and prey may have differential temporal activity patterns, the covariance between interacting species’ patterns (i.e., temporal synchrony) will structure species interactions through time. Synchronized activity timing, for example, can increase predator attack rates on prey (Dou et al. 2019), while asynchronized activity would be expected to reduce predator-prey interactions. In this section, we first provide previously published empirical examples of how predator-prey activity patterns manifest in nature before using theory to begin to explore the consequences of activity mean, variation, and timing for stability.

### Empirical examples

A non-exhaustive literature search using the terms [forage, active, time, predator, prey] located studies that compared predator and prey activity at the daily, lunar, and seasonal scales. These studies overwhelmingly focused on activity timing between predators and prey and provide evidence that both synchrony and asynchrony in activity timing are possible (table S1). At the daily scale, synchronized activity was reported between predator-prey pairs including wolves and moose (Vander Vennen et al. 2016), tigers and wild boar (Dou et al. 2019), and coopers hawks and small birds (Roth and Lima 2007), which were all active at the same time of day (table S1). At the lunar and seasonal scale, synchronized examples include owls and mice both increasing activity during the new moon (Clarke 1983), and wolves and moose both remaining active all year (Ditmer et al. 2018). Synchronized activity timing, on average, would be expected to increase encounters and predator attack rates on prey. For example, tigers preferentially consumed prey (wild boar) that were lower in density but active at the same time (Dou et al. 2019).

Asynchronized activity was also observed, which should on average reduce predator-prey encounters during mismatched activity times while also potentially focusing attack rates on any periods of overlap (e.g., dusk, dawn). For example, activity in nocturnal snow leopards was asynchronized with diurnal ibex but attacks still occurred during the crepuscular period when light conditions appeared to improve prey catchability (Johansson et al. 2022). At the lunar and seasonal scale, asynchrony arose when a predator increased activity during a different lunar phase (as in jaguar and sea turtle; Carrillo et al. 2009) or different season (hornets and honeybees; Monceau et al. 2013) than prey, or when the predator did not respond to the lunar phase, but the prey did (as in ocelots and armadillos; Pratas-Santiago et al. 2016). Several other examples indicated that a predator might be asynchronized with one prey but synchronized with another prey (table S1). For example, nocturnal pine marten are synchronized with nocturnal wood mouse but asynchronized with diurnal squirrels (Caravaggi et al. 2018).

Although the variation in activity was not directly quantified, predators and prey in most examples were nocturnal, diurnal, or crepuscular and could therefore be considered to have variable activity patterns at the daily scale relative to cathemeral species that can be considered more constantly active. At the seasonal scale, constant activity was reported in most studies, likely due to researchers studying seasonal interactions in predators and prey that remain active during the studied seasons (table S1).

Mean activity was also not directly reported in the sample studies, but many of the examples appear to be ‘faster’ predators and prey (e.g., ibex and snow leopard (Johansson et al. 2022), barracuda and red snapper (Rooker et al. 2018), lynx and hare (Shiratsuru et al. 2023); table S1). Interestingly, these ‘faster’ species with higher mean activity should have higher metabolic and growth rates, along with a suite of other morphological, physiological and life history differences, compared to ‘slower’ species as per the pace-of-life syndrome hypothesis (Ricklefs and Wikelski 2002). For example, both lynx and hare have high mean activity rates and are also known to have high metabolic rates and strong interactions leading to high amplitude population cycles in both species (Menzies et al. 2022, Shiratsuru et al. 2023). This suggests that high mean activity in both predators and prey potentially leads to strong interactions that are known to reduce stability (McCann et al. 1998). However, the influence of predator and prey activity patterns on interaction strengths and stability has not yet been widely studied.

### Time-varying predator-prey theory: model and implications

#### i) The model

In the section above, we demonstrated that organisms differ widely in activity mean, variation, and timing. Towards motivating further theoretical development in this important area, we seek to begin to highlight the potential implications of the activity mean, variance, and timing on one of the fundamental building blocks of food webs, predator-prey interactions. In box.1 below we introduce in more detail a simple predator-prey model where each species’ activity is periodically-forced to create differential temporal activity patterns that align with our activity traits discussed above. Here, we follow the well discussed theoretical assumption that attack rates scale with the mean relative speed of interacting organisms (hereafter, **the mean relative speed** (**MRS**) assumption). Theory developed around the movement of two different particles (e.g., predator and prey) has shown that, all else equal, the mean relative speed of the two types of particles predicts the rate of collision (i.e., predator and prey attack rate) within a given area (Gurarie and Ovaskainen 2013, DeLong 2021).

With this well used assumption in tow, we theoretically consider two biologically plausible activity/attack rate assumptions. First, we assume that the encounter or attack rate is entirely set by the mean relative speed assumption (hereafter **CASE 1: MRS** see box. 1). This suggests that attack rates will be maximized when both predator and prey are moving. Note, that the MRS assumption includes predators employing sit and wait foraging as they can still consume if the prey alone is moving. Second, still within this MRS framework, we then consider the possibility of prey using refugia to avoid predation. As a first pass, we do so by incorporating the added assumption that prey may be using refugia when they are less active (i.e., when moving less than their average activity rate). Here, when prey are more stationary, their presence in (or near) refugia ought to limit encounter rates, potentially below what would be expected by the MRS alone. In this case, we assume attack rates still follow the MRS assumption when prey are active (prey activity > average activity), however when prey are less active (prey activity < average activity), we then assume that the attack rate is then set solely by the prey movement alone (**CASE 2: MRS + Refugia** see box. 1). Here, prey are still accessible to predators when moderately active, but a zero movement of prey, for example, assumes they are fully hidden in the refugia and so experience zero encounter rates regardless of predator movement rates.

Finally, to explore a range of dynamical outcomes and to incorporate potential differences in mean activity rates, we consider two baseline scenarios for both cases above: i) **a slow mean activity** (i.e., weak attack rate/interaction strength) scenario; and ii) **a fast mean activity** (i.e., strong attack rate/interaction strength) scenario. In the Rosenzweig-MacArthur predator-prey model used here as a theoretical foundation, increasing attack rates are known to incite a transition from stable, monotonic dynamics through unstable, oscillatory dynamics (referred to as the “paradox of interaction strength/energy flux” by Rip and McCann (2011)). Thus, by accounting for different mean activity rates (i.e., attack rates following the MRS), these scenarios also allow us to explore the effects of variable activity within different dynamical regimes.

Importantly, these two regimes have been frequently identified as producing alternative responses to changes in attack rates (interaction strengths). Increasing attack rates in weak, non-excitable equilibrium dynamics tends to increase stability (e.g., called mean-driven stabilization), as it moves mean predator away from near zero densities, decreases return time, and reduces CV in stochastic cases (Gellner et al. 2016, McCann et al. 2021). In contrast, strengthening already strong oscillatory interactions further destabilizes predator-prey dynamics (i.e., called variance-driven instability), as it promotes overconsumption, thus increasing oscillations and CV and causing near-zero densities during cycles (McCann et al. 2021).

Altogether, we explore four model settings (2 cases x 2 baseline scenarios), within which we investigate the implications of activity variation and timing. That is, in each of these model settings, we explore a range of activity cycle amplitudes and alter the synchrony between predator and prey activity patterns, from perfectly synchronized to completely asynchronized in time.

###### Box 1: Theoretical model and simulation

To examine how the mean, variance, and timing of different activity strategies may influence the stability of predator-prey dynamics, we use a predator-prey (consumer-resource) model, with the addition of sinusoidal functions for the predator and prey’s activity rates, such that

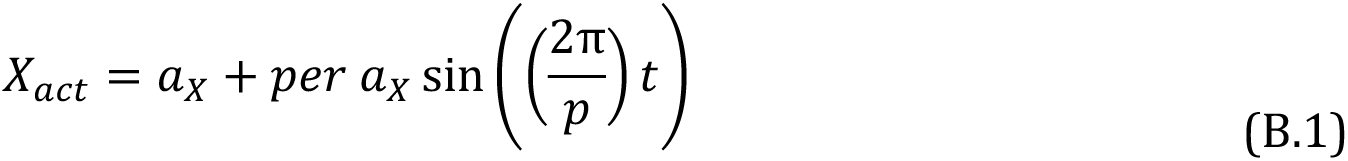

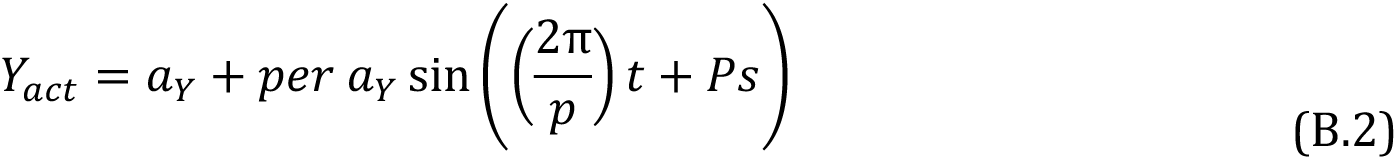

where *X_act_* and *Y_act_* are the sinusoidal functions of the prey and predator’s activity respectively, *a_i_* is the mean activity rate, for species *i*, *per* is the scalar percentage that allows us to toggle the activity variation for the predator and prey, *p* is the forcing frequency, and *Ps* is the phase shift where we can alter the synchrony between predator and prey. With these two functions, we force the attack rate parameter, 𝛼_+,_(𝑡), with our first assumption, the **mean relative speed (MRS)** by

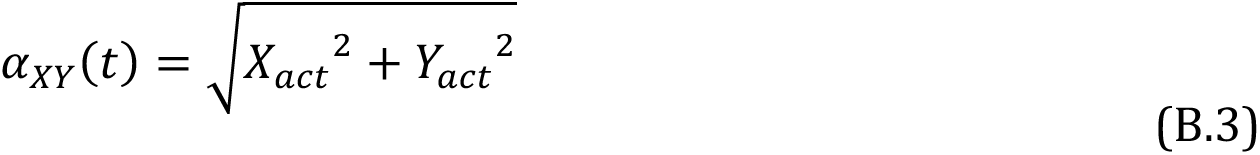

where the attack rate varies such that 𝛼*_XY_*(𝑡) is a function of the mean relative speed. Here, the MRS assumption predicts the collision rate (i.e., attack rate) of two moving particles (i.e., predator and prey) within a given area (Gurarie and Ovaskainen 2013, DeLong 2021).

For our second case, **MRS + Refugia**, we force the attack rate parameter, 𝛼*_XY_*(𝑡), by

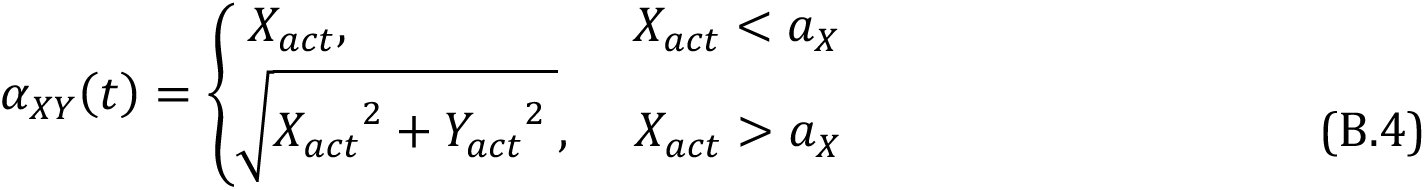

where the attack rate now varies such that 𝛼*_XY_*(𝑡) is a function of the mean relative speed with refugia. Here, prey use refugia to avoid predation when they are less active (prey activity (*X_act_*) < average activity (*a_X_*)) where the attack rate, 𝛼*_XY_*(𝑡), solely depends on the prey’s movement, however, when the prey are more active (prey activity (*X_act_*) > average activity (*a_X_*)), the attack rate, 𝛼*_XY_*(𝑡), follows our mean relative speed assumption.

Next, using the Rosenzweig-MacArthur predator prey model with a Type II functional response (Rosenzweig and MacArthur 1963), the attack rate function that scales with activity is substituted into the model, giving

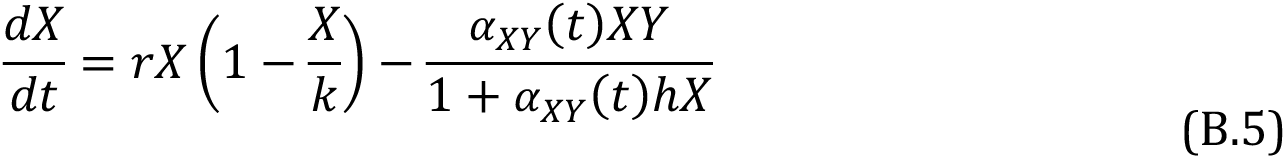

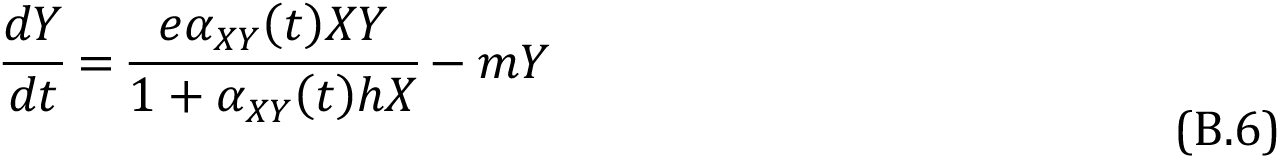

where *X* and *Y* are the respective prey and predator state variables, *r* is prey growth rate, *k* is prey carrying capacity, *h* is the handling time, *e* is conversion efficiency, and *m* is the predator’s mortality. Finally, 𝛼*_XY_*(𝑡) is the predator’s attack rate on the prey which is a function of time proportional to the sinusoidally forced predator’s activity pattern, resulting in temporal patterns emerging through these parameters.

From the above model, the predator’s coefficient of variance (CV) is calculated from numerical simulations of two different cases where the mean activity rate of both the prey and predator are either slow (*a_i_* = 0.25) or fast (*a_i_* = 0.7). Numerical simulations were tracked over the interval range of 900 to 1000 time-units to minimize transient influence, and the CV was computed over this range. Other parametric values can be found in the figure captions. Within each case, the timing and variability of both the prey and predator’s activity strategies are altered to determine the overall effect on interaction strengths and stability. All numerical simulations are coded in Mathematica 12.0.0.

#### **ii)** Results

The above model is a forced predator-prey model whereby the attack rate parameter is a time-varying function of the sinusoidally forced predator and prey’s activity (box. 1). Here, we consider variability in the dynamics (measured as the coefficient of variation – CV – of the predator, where CV = standard deviation (SD)/mean (μ)) as an indicator of stability that is aligned with our empirical examples/results above (i.e., small mouth bass and lake trout change in their variability (CV) across different timescales; figure 3f). As such, there are two main ways that dynamics become less stable in this model: by increasing variance (SD) or by decreasing mean predator density (*μ)*. Clearly, deterministic predator-prey cycles make dynamics more variable, regardless of external forcing in activity. However, fluctuating attack rates may change the average attack rates experienced, which in turn may alter both the mean densities and the underlying stability of our model. Moreover, introducing an additional oscillator into the system could further change the variability and frequency of these dynamics (see Supplementary Material S2). Here, we explore these interactions.

Across the activity-driven attack rate cases and baseline scenarios explored here, our predator-prey dynamics may thus become more variable for multiple reasons. First, the temporal interaction of predator and prey activity may alter the mean attack and given that the mean attack rate is known to alter stability, this change alone can impact dynamics (e.g., strong attack rates yield cycles; Rip and McCann 2011). As an example, if predator and prey activity is out of phase, we may see a massive decline in attack rates and much less oscillatory dynamics. Second, forced consumer-resource (predator-prey) models can have variable dynamics simply from the periodic forcing itself (here, driven by time varying activity/attack rates). For example, given parameters that yield a stable equilibrium, externally forced dynamics are known to generally produce periodic predator-prey densities around the deterministic equilibrium – such that the magnitude of the variation scales with the amplitude of the forcing (Bieg et al. 2023). On the other hand, the forcing of an already oscillating predator-prey dynamic is more complicated as variability in the dynamics represents the interaction between the deterministic predator-prey cycles and the periodic forcing, which is importantly dependent on the frequency of forcing (Bieg et al. 2023). Because forcing and changes in predator–prey cycles both affect variability (SD and thus CV), we therefore assess how altered activity patterns influence these two mechanisms and overall stability (CV). Finally, with respect to the periodic forcing mechanism, the frequency of the forcing also potentially impacts the dynamics and stability. Slow forcing, for example, can be tracked by a population (thus driving variation in dynamics discussed above) but fast forcing is perceived by the organism more as an “averaging” effect, thus reducing the impacts driven by periodic forcing.

### CASE 1: Mean relative speed (MRS) assumption (figure 5A)

To explore this initial scaling relationship between activity and encounter rates, we first demonstrate in figure 5A how temporally-forced activity of predator and prey (grey solid and dashed lines, respectively) translate to the time-varying attack rate under the MRS assumption (thick-black solid curve). We do so for both synchronous (figure 5Ai) and asynchronous (figure 5Aii) predator and prey activity timing when both mean activity rates are slow (a=0.25; figure 5Aiii) and fast (a=0.7; figure 5Aiv, S3.4). We note immediately that asynchronous behavior reduces the amplitude of the time varying attack rates (figure 5Aii compared to figure 5Ai), while doubling the periodic attack rate’s frequency (Supplementary Material S2.1). Further, we see that under the MRS assumption the mean attack rate increases from 0.35 to 0.43 (see figure 5Ai and figure 5Aii insets respectively) as the behavioral asynchrony now elevates the mean attack rate (figure 5Aii).

**Figure 5:**
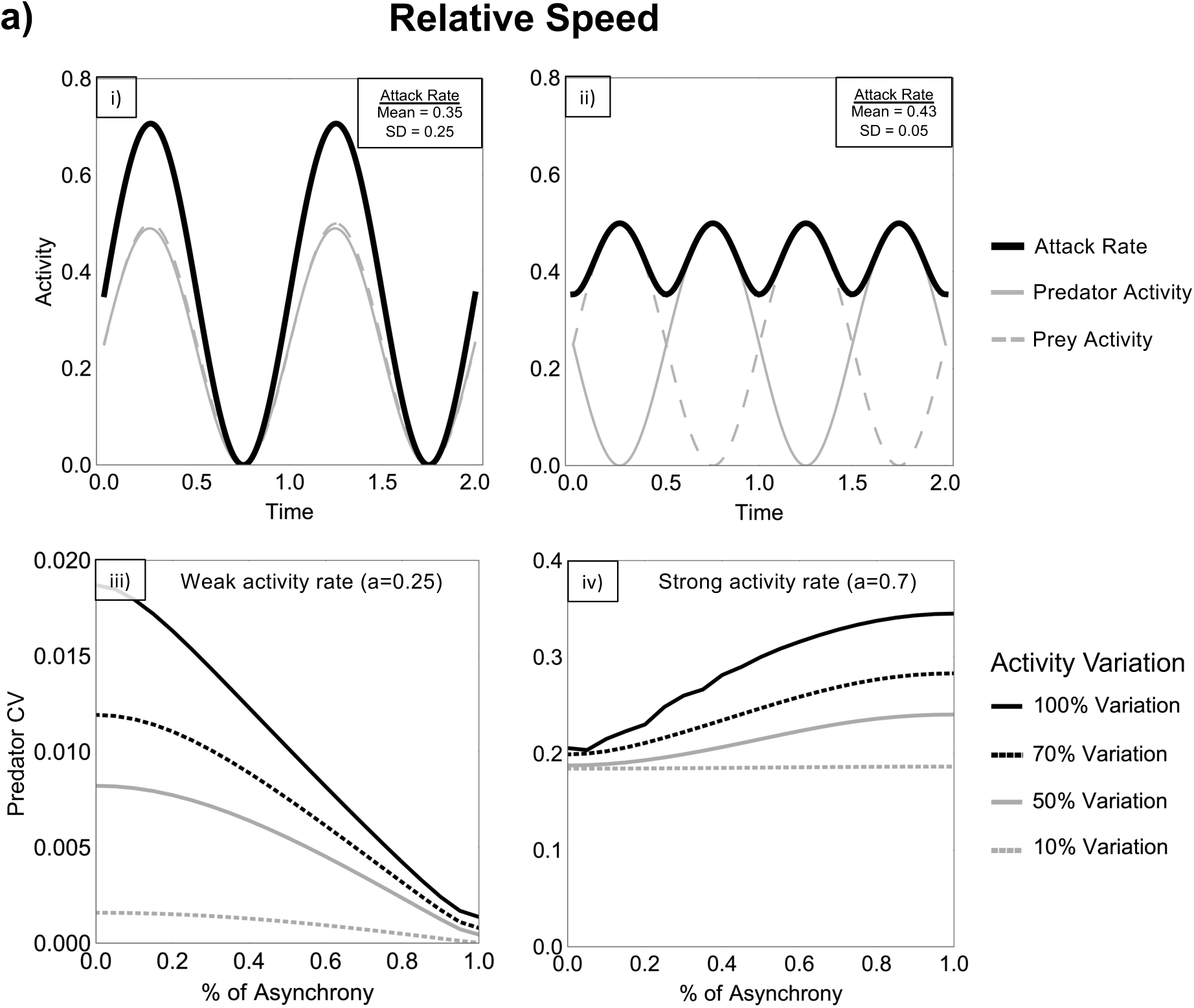
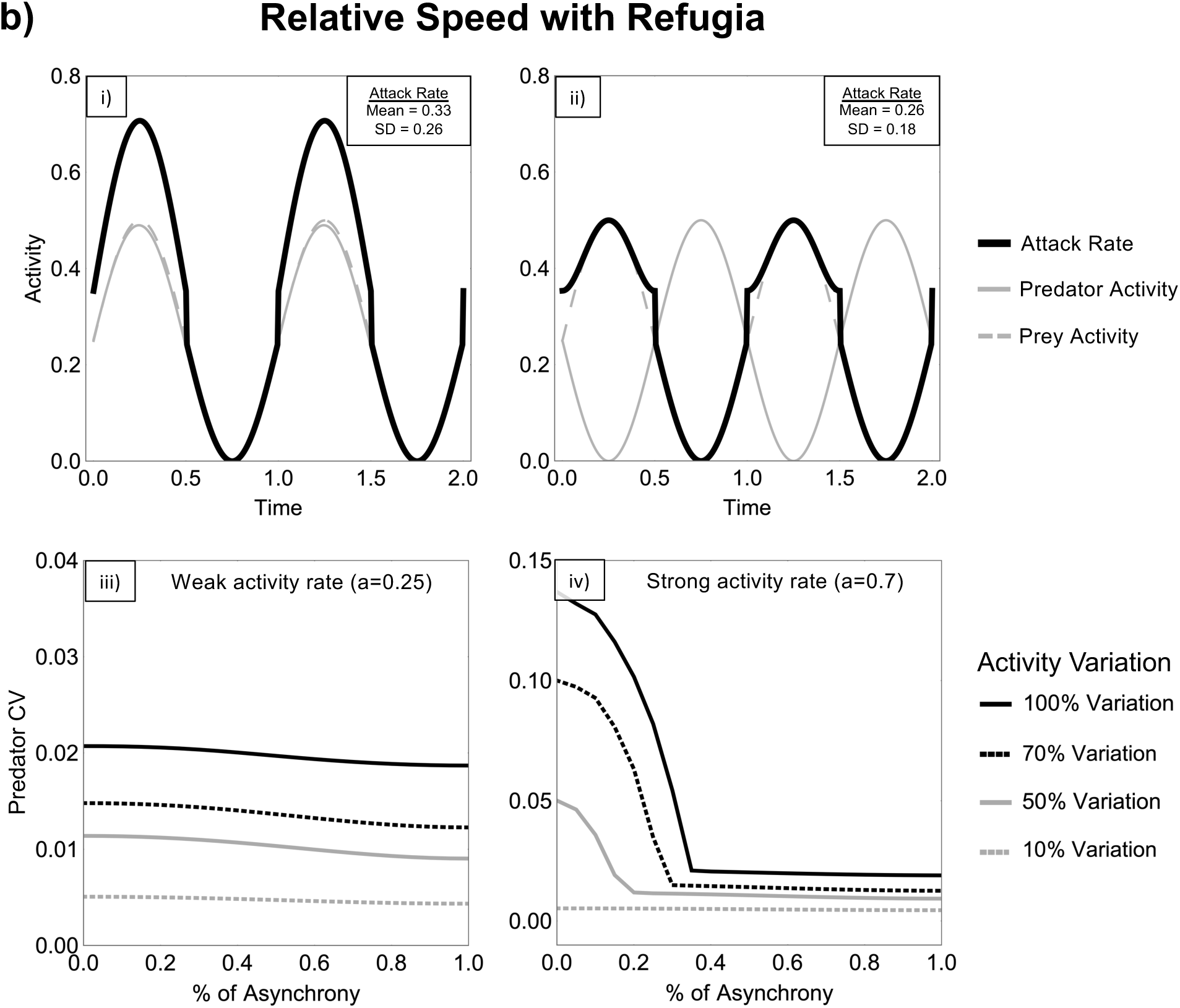
Panel A: Relative maximum attack rate scales with the activity patterns of the predator and prey when the activity patterns are synchronous (i: *Ps* = 0) and asynchronous (ii: *Ps* = 1), and the predator’s coefficient of variation (CV) as these activity patterns between predator and prey become more asynchronous and variable when the mean activity rate is slow (iii: a=0.25) and fast (iv: a=0.7). See figure S3.4 for the scaling of the maximum attack rate when the activity rate is strong. Activity variation is the scaler percentage (*per*) of the predator and prey’s activity in panel i and ii. Parametric values: *r* = 2.5, *k* = 2.0, *m* = 0.2, *h* = 1, *e* = 0.7, *p* = 1. **Panel B:** Relative maximum attack rate with refugia – scales with the activity patterns of the predator and prey when the activity patterns are synchronous (i: *Ps* = 0) and asynchronous (ii: *Ps* = 1), and the predator’s coefficient of variation (CV) as these activity patterns between predator and prey become more asynchronous and variable when the mean activity rate is slow (iii: a=0.25) and fast (iv: a=0.7). See figure S3.5 for the scaling of the maximum attack rate when the activity rate is strong. Activity variation is the scaler percentage (*per*) of the predator and prey’s activity in panel i and ii. Parametric values: *r* = 2.5, *k* = 2.0, *m* = 0.2, *h* = 1, *e* = 0.7, *p* = 1.

Given the two mechanisms (forcing and attack rate) impacting stability (CV) discussed above, we first examine the effect of forcing given a slow mean activity rate (i.e., weak attack rate; figure 5Aiii), where the underlying deterministic model (i.e., no forcing) is stable. Here, predator variance clearly depends on the magnitude of forcing imposed (i.e., activity variation), as stronger forcing results in more variable dynamics (see grey dashed to solid black lines in figure 5Aiii representing changing amplitude of forcing relative to the maximum attack rate shown in panels i and ii). However, we also see that this result is dependent on the synchrony of predator and prey activity timing. From above, we expect the mean predator density to increase (relative to prey density) with asynchrony as the mean attack rate increases – based on a well-known result from predator-prey (consumer-resource) theory that increasing attack rate drives a weakly interacting predator to higher mean densities (thus decreasing its CV as CV=SD/*μ;* figure 5Aiii, S3.6a). This has been referred to in the literature as mean-driven stabilization (McCann et al. 2021) as it moves the predator equilibrium away from collapse (i.e., away from near zero densities). Second, as asynchrony increases, the attack rate variance – due to the amplitude of the periodic forcing of both species – decreases from 0.25 to 0.05 (see figure 5Ai and figure 5Aii insets respectively), suggesting that the variance in predator dynamics due to forcing should also decrease (decreasing CV as the SD also decreases; figure S3.6b) – known as variance-driven stabilization (McCann et al. 2021). Additionally, the periodic attack rate’s frequency doubles, thus exerting a stronger pull towards the mean equilibrium on the predator-prey dynamics (Supplementary Material S2.1). Thus, all these mechanisms suggest stabilization, and this is precisely what happens (figure 5Aiii, Supplementary Material 2.1). In other words, given some amount of activity variation and a weakly-coupled predator-prey interaction, asynchrony in predator and prey activity patterns increases stability (via decreased CV), with the magnitude of this stabilization result dependent on the variance in activity (figure 5Aiii).

For the fast mean activity (strong interaction; figure S3.4) scenario, we see that the system becomes less stable (predator CV increases; figure 5Aiv) as the activity patterns become more asynchronous (relative to an already unstable underlying model). In this case, the two mechanisms (i.e., forcing and attack rate) work in the opposite direction. Asynchrony in the periodic forcing function (which decreases the maximum attack rate’s amplitude) decreases the variance of the predator’s periodic attack rate from 0.7 to 0.14 while doubling its frequency (see figure S3.4i and S3.4ii insets respectively; Supplementary Material S2.1) -- both suggest that we should see a decrease in variation in population dynamics from forcing. At the same time, the mean attack rate increases from 0.99 to 1.20 (see figure S3.4i and S3.4ii insets respectively). This is well known from predator-prey (consumer-resource) theory to ramp up the size of the predator-prey cycles (Rip and McCann 2011). While the increase in mean attack rate outweighs the dampening effect of the reduced forcing amplitude – driving higher CV and reducing stability (figure 5Aiv) – our supplementary analysis further shows how changes in the average attack rate, variability, and frequency alter the predator-prey dynamics across the phase-plane, thus reinforcing this destabilization (Supplementary Material S2.1; figure S2.1).

In summary, under the MRS assumption, there are clear and strong stabilization and destabilization effects in response to mean activity rates (weak and strong attack rates), variance in activity (variance in attack rates) and the timing of activity patterns (degree of asynchrony between predator and prey activity) – all three of our activity traits discussed above. Under the well-known MRS assumption, dynamics and food web resilience are therefore strongly influenced by activity/attack rate temporal patterning.

### CASE 2: Mean relative speed with refugia (MRS + Refugia) assumption (figure 5B)

As above, we first show how this assumption takes the activity of the predator and prey in time (grey solid and dashed lines respectively) and determines the maximum attack rates (thick-black solid curve) when predator and prey activity patterns are synchronous (figure 5Bi) and asynchronous (figure 5Bii). With the assumption of refugia added we note immediately that asynchronous behavior again reduces the amplitude of the time varying maximum attack rate, particularly on the “high” end of the fluctuations; note the pronounced skew to low values in the resulting attack rate curve (figure 5Bii; Supplementary Material S2.2). However, the periodic attack rate’s frequency remains the same when activity patterns are asynchronous compared to when they are synchronous. We finally note that under the refugia assumption, the mean attack rate now decreases significantly as predators and prey become more asynchronous in their patterns (see mean attack rate = 0.33 to 0.26; figure 5Bi and figure 5Bii insets respectively). This occurs as the slower prey speeds imply refugia and thus produce weaker attack rates than produced in the MRS assumption (figure 5Aii).

As above, we first examine the response of the slow mean activity rate (weaker attack rate; figure 5Biii). In contrast to the MRS case above, we now expect the mean predator density to decrease with asynchrony as the mean attack rate decreases (figure S3.7a), based on predator-prey theory that suggests a weakly interacting predator will be driven to dangerously low densities (thus increasing CV) as attack rates (i.e., consumption) decrease (McCann et al. 1998). On the other hand, the variance (amplitude) in the forced attack rate also decreases with asynchrony (see figure 5Bii; Supplementary Material S2.2) and therefore ought to decrease predator variance (decreasing CV; figure S3.7b). Thus, one mechanism (weaker mean attack rate) suggests destabilization (increasing CV) and the other mechanism (reduced forcing amplitude) suggests stabilization (decreasing CV) – two mechanisms working against each other. Here, we see that these mechanisms indeed counter each other and create a very weak stabilization effect with synchrony (figure 5Biii, Supplementary Material 2.2). Again, the amplitude of forcing (i.e., activity variance) intuitively scales this weak CV response.

Finally, with the fast mean activity (strong attack rate/interaction strength) scenario, we find that asynchrony has a stabilizing effect on predator-prey dynamics with this added refugia assumption (figure 5Biv) – opposite of the result we found above for the MRS assumption with fast mean activity/strong attack rates. Here, the two mechanisms work together to drive this stabilization. The decreasing amplitude of our periodic forcing function with increasing asynchrony should decrease the variance of the predator’s attack rate (figure 5Bii), and the decreased mean attack rate now mutes the size of the predator-prey cycles, eventually removing them if the attack rate decreases enough to cross the Hopf bifurcation (as seen in figure S2.2B1, S2.2B2). Accordingly, asynchrony stabilizes this otherwise oscillatory system by jointly reducing variability and damping predator–prey cycles, as illustrated by shifts in phase-plane structure and equilibria in Supplementary Material 2.2.

In summary, under the refugia assumption, the implications of the activity metrics (mean, variance, and timing) on predator-prey dynamics are considerably different than the well-known MRS result but nonetheless shows that time varying activity rates likely have enormous implications for dynamics and stability. We would argue that understanding and further developing the link between activity and attack rates is fundamental to understanding predator-prey interactions and thus food web dynamics. The MRS assumption although theoretically reasonable, may be biologically unreasonable when we think about the role of space and behavior. Further, the ability to map activity behavior in detail now puts us in a position to empirically harness this data towards understanding its role in predator-prey and ultimately food web dynamics and stability.

### Conclusions and future directions

Here, we demonstrate diversity in the timing, mean, and variation in activity among taxa and between predators and prey that arises at daily, lunar, and seasonal timescales. Importantly, both synchronized and asynchronized activity timing was reported between predators and prey in the literature. Based on our theoretical models, differences in activity synchrony have the potential for large consequences for stability that depend on the mean, variation, and timing of predator and prey activity. We explored two cases, the first case where activity rates map to attack rates following the mean relative speed of predator and prey (defines collision rate of random movement) and a second case that assumes prey use refugia when they are moving slower than their average mean speed. These two cases highlight different stability responses suggesting a critical need for empirically developing our understanding of the relationship(s) between activity rates and attack rates. Below, we conclude by discussing avenues for future directions based on our findings.

### Attack rate and activity relationship

These two cases presented here represent alternative assumptions about how activity influences attack rates. The baseline MRS assumption is a longstanding theoretical construct and likely valid in many empirical cases. For example, many predators must move to locate and consume prey, which should generate positive relationships between activity and attack rates. Prey consumption by predatory lionfish was shown to peak with maximum lionfish activity rates and reach a minimum when lionfish are least active (Green et al. 2021). The movement rate of predatory wolves also predicted attack rates on moose (Vander Vennen et al. 2016). We also modified this assumption to account for refugia (MRS+Refugia), which should hold when prey experience lower predation risk when inactive (e.g., by using refugia) and restrict predation events to times when prey are active. For example, the timing of activity in small birds did not correspond with the timing of intense predatory activity of Cooper’s hawks, suggesting the use of refugia to avoid predation (Roth and Lima 2007). Abiotic conditions like light or temperature that place constraints on predator performance might also underpin the correlation between activity and attack rates. For example, darkness is thought to limit attack rates by snow leopards on inactive ibex during nocturnal periods (Johansson et al. 2022).

While both assumptions are reasonable and have empirical and theoretical backing, there are numerous other ways activity and attack rates may scale. For hares that rest during the day in the open with no refugia, attacks by more consistently active (cathemeral) Canadian lynx were equally likely during the day or night (i.e., regardless of prey activity timing; Shiratsuru et al. 2023). Invasive hornets increased attack rates on native honeybees at midday, not because of changes in honeybee activity, but because of improved predator performance under warmer temperatures (Monceau et al. 2013). In these examples, the activity of prey does not play a role in the attack rate. Finally, activity associated with behaviors other than foraging (e.g., reproduction, dispersal, thermoregulation) could periodically weaken or even change the direction of the correlation between activity and attack rate. More work is required to quantify the many possible ways that activity could relate to attack rate. However, the requirement for many animals to move to find and consume resources suggests that activity and attack rate can be positively related across many diverse taxa.

### Predator-prey activity across temporal scales

We have considered how diverse activity patterns arise among different taxa and might influence predator-prey interactions. However, activity is not a static trait and can vary at multiple temporal scales (Helm et al. 2017). Our literature search uncovered examples of daily activity patterns that shift across lunar, and seasonal timescales (table S1). For example, the timing of daily activity changes from crepuscular in summer to diurnal in winter in moose and wolves (Ditmer et al. (2018); also see table S1 for more examples of monthly and seasonal shifts in the timing of daily activity). Many examples also exist of facultative cathemerality in mammals, where the extent of cathemeral activity (i.e., foraging in both day and night) changes during certain times of the lunar or seasonal cycle (Cox and Gaston 2024). In some cases, predator-prey pairs are synchronized on one scale but asynchronized on different scales. For example, wildcats and their rabbit prey are both most active at night during the winter (i.e., synchrony), but during the remainder of the year, rabbits switch to daytime activity (i.e., asynchrony) (Martín-Díaz et al. 2018). For simplicity, our theoretical models focused on attack rate forcing at relatively small timescales (e.g., predator-prey periodicity >> forcing periodicity). While beyond the scope of this paper, a scale dependent theory is required to fully understand the behavioral implications of abiotic forcing and activity patterns on stability.

### Flexibility in activity as an adaptive response

In addition to variation in activity patterns at multiple timescales, activity patterns could also shift through an organism’s lifetime, through ontogenetic shifts and responses to external stimuli, like anthropogenic activities (Gilbert et al. 2023, Cox and Gaston 2024) or in response to spatial variation in habitat or ecosystem characteristics. For example, a tropical reef fish (*Siganus lineatus*) was diurnal along the shoreline but possessed the flexibility to switch to nocturnal foraging on reefs (Fox and Bellwood 2011). Lake trout were also shown to spend more of the day active when inhabiting lakes with smaller vs. larger prey (Blanchfield et al. 2023). Additional responses to changing biotic conditions includes lionfish (Benkwitt 2016) and Tasmanian Devils (Cunningham et al. 2019) shifting the duration and timing of their daily activity as intraspecific densities increased.

In a similar way to diet and habitat switching, migration and physiological adaptations, flexible shifts in the timing or magnitude of activity and rest could be an additional mechanism allowing organisms to couple and decouple from fluctuating conditions in space and time. For example, the capacity for activity during day or night, could increase flexibility and better allow an organism to avoid unfavorable conditions and take advantage of favorable periods, although performance might be lower compared to species that are strictly active in the day or night (Cox and Gaston 2024, Bloch et al. 2013). More work is required to explore the underlying mechanisms shaping activity flexibility across taxa (e.g., endotherms vs. ectotherms) and to connect activity flexibility with consequences for organismal performance, population dynamics and food web stability in a variable world. Answering these questions are especially pressing given the recent recognition that activity likely plays a role in shaping sensitivity to anthropogenic impacts like warmer temperatures and increased light pollution (Gilbert et al. 2023, Cox and Gaston 2024).

### Activity rates and temporal compartmentation

From a food web perspective, variation in activity rates and timing suggest the potential for activity driven temporal compartments in food webs akin to the spatial compartments that impact stability (McCann et al. 2005, McMeans et al. 2015). Specifically, we can define temporal guilds as species that show synchronized dramatic highs and lows in activity (i.e., switching on and off). For example, at the seasonal scale, both aquatic (McMeans et al. 2015, 2020) and terrestrial food webs support species that maintain their activity all year vs. those that reduce activity during harsh winter seasons (Humphries et al. 2017). These divergent activity patterns and different temporal guilds (e.g., day versus night, summer/wet vs. winter/dry, etc.) effectively create temporal food web compartments. Here, the analogue to spatial compartments that have habitat generalists coupling food webs in space are temporal generalists like cathemeral and winter-active species that couple night-day and summer-winter webs for example by being generally active at the given timescale (Tattersall 2008, Humphries et al. 2017). While not well explored, this temporal compartmentation may be expected to play a key role in stability akin to the potent role spatial generalists play in spatial food web theory. Nonetheless, more synthetic empirical and theoretical work with respect to these compartments are required and even demanded given the data emerging on activity patterns. Finally, this work will naturally tie into the intersection of temporal activity and space, an area that we ignored here but will be critical to fully understanding the role of activity in species interactions, food webs and ecosystems.

## Author Contributions

EKS and BCM contributed to the conceptualization of the study. AMS, CB, KSM and BCM contributed to literature investigation and analysis. AMS, BMS contributed to the method analysis. AMS, CB and KMS contributed to the theoretical analysis. AMS, CB, KMS and BCM contributed to visualization and figures. AMS led the final draft preparation and submission stages with comments from all authors being received prior to submissions.

## Biographical Narrative

Alexa Scott is a PhD candidate, Kevin McCann is a professor, and Carling Bieg is an assistant professor at the University of Guelph in Guelph, Ontario, Canada. Emily Studd is an assistant professor at Thompson Rivers University in Kamloops, British Columbia, Canada. Brett Studden is PhD candidate at the University of Toronto Mississauga, and University of Toronto in Toronto, Ontario, Canada. And Bailey McMeans is a professor at the University of Toronto Mississauga in Mississauga, Ontario, Canada.

## Data and Code Accessibility Statement

Raw abiotic data are publicly available through the Historical Climate Data database from Environment and Climate Change Canada and via the suncalc package in RStudio, as described in our methods section and table S2. A subset of the activity data is publicly available through Movebank, as detailed in our methods section and table S2; additional activity data are not publicly available. Refer to the methods and table S3 for further filtration of the raw data used for the manuscript analysis. All code used to reproduce the analyses and figures has been archived in a Zenodo repository (version 1.0).

