## Supplementary Material for "Activity Patterns Structure Food Web Interactions Through Time"

### S1 Methods

#### S1.1 Abiotic Conditions

Temperature and snow depth data were gathered from Algonquin Park, East Gate, Ontario, Canada (45.53°N, -78.27°W) through the Historical Climate Data database from Environment and Climate Change Canada (table S2). Both solar irradiance and moon illumination data were calculated using the suncalc function in RStudio v. 2023.6.0.421 with the same coordinate position (table S2). This location, and the dates selected for examination (table S3), were chosen as they best resemble the environment for which the exemplar species, *Micropterus dolomieu*, occupies. All abiotic conditions were standardized in RStudio v. 2023.6.0.421 for comparison.

#### S1.2 Activity

We gathered acoustic telemetry activity data for smallmouth bass (*Micropterus dolomieu*) and lake trout (*Salvelinus namaycush*) using a pre-existing Vemco Positioning System installed in Smoke Lake (45.5172°N, -78.6816°W; Algonquin Provincial Park, Ontario, Canada). We also gathered satellite derived activity data from Movebank (movebank.org) for black-backed jackal (*Canis mesomelas*: study name "Black-backed jackal, Etosha National Park, Namibia") and African elephant (*Loxodonta africana*: study name "African elephants in Etosha National Park (data from Tsalyuk et al. 2018)") from Etosha National Park (19°S,16°E), Namibia, Africa, and for mallard duck (*Anas platyrhynchos*: study name "LifeTrack Ducks Lake Constance") and great cormorant (*Phalacrocorax carbo*: study name: "Great Cormorant Lake Constance MPIAB") in Lake Constance, Baden-Württemberg, Germany (table S2). Two organisms from three different ecosystems (i.e., aquatic, terrestrial, aerial) were chosen to highlight the diverse variety of activity patterns among species but also within different ecosystems. This selection was also restricted by sufficient resolution quality within the dataset (≤ 1hr fix intervals) and duration of the dataset (continuous data collection for ≥ 1 year) to allow for analysis across the daily, lunar, and seasonal timescales.

S1.3 Movement Analysis:

The acoustic telemetry datasets that came from Algonquin Provincial Park are part of an ongoing study at the Harkness Laboratory of Fisheries Research of the Ontario Ministry of Natural Science. Detection data from tagged lake trout and smallmouth bass (V13AP, V9AP, and V9P tags) were collected during three downloads (September 2021, May 2022, and June 2023). Positional data was computed using INNOVASEA's proprietary hyperbolic positioning algorithm and data were filtered to remove positions outside of the lake perimeter, impossible movements and dead or missing fish. Additionally, a time-elapsed greater than or equal to 28 mins between consecutive positions was removed to reduce a higher error potential when calculating activity rate (table S5). Finally, system error (HPEm) was determined using Bayesian quantile regression between HPE and HPEm for 6 reference tags across months and positions with a median error greater than 15 m were removed.

The raw data from the Movebank Data Repository is formatted as comma-separated values (CSV) files. Each file typically includes data from all individuals within a particular study, with each individual assigned a unique identifier (tag) to differentiate them. Although the source and structure of the data may vary depending on the specific study, several common variables are present across datasets. Using timestamp and locational coordinates, the Haversine formula was applied to calculate speed (m/s) for each individual. This formula determines speed by dividing the distance traveled (in meters) by the time difference between consecutive fixed points (in seconds). The Haversine formula was chosen because it accurately measures distances between two points on a curved surface, and has been used in previous animal movement studies (Curry 2014, Santhosh and Idicula 2017). All datasets were then filtered with a resolution ≤ 1hr to achieve consistent fix rates for comparison across the daily timescale and to avoid significant data loss for accurate comparisons (see table S5 for filtration information). Since the telemetry data resolution was already pre-filtered with a max time difference of 28min between each position, the following analyses were rerun at resolutions ≤ 28min intervals, see supplementary material Fig S8, S9, S10.

To gain an understanding of how these different activity patterns may correspond with one another across and within each ecosystem, 2D velocity was measured across the daily, lunar, and seasonal timescales for all species. For each ecosystem, the month during which both species exhibited relatively high activity was selected for daily-scale comparisons. When peak activity periods differed between species, the month in which both displayed the highest overall activity was used (July for aquatic, April for terrestrial). June was chosen for both aerial species, the month when they are both relatively inactive to examine their ecosystem foraging behaviour and avoid migratory behaviour (Dingle and Drake 2007). Average hourly velocity was measured for the daily scale across the entire month to avoid any possible noise, and if multiple years were available within the dataset, all days within that month across all available years were used in the analysis (table S3). For the lunar timescale, 30-day periods corresponding to a full lunar cycle (from full moon to new moon and back to full moon) were analysed for the periods when each species exhibited the highest average daily activity. If multiple lunar cycles of high activity periods were available, the daily average velocity was taken from all cycles to gain the most accurate representation of the species behaviour and activity pattern (table S3). Finally, for the seasonal timescale, monthly average velocity was calculated using the entire dataset (table S3).

After the 2D velocity was calculated for each species across each timescale, the mean, standard deviation (SD), and the coefficient of variance (CV) of activity were calculated (figure 3, S3.2, S3.3). To examine activity synchrony, activity rates were classified relative to the mean movement rate using a t-based threshold defined as mean ± $t$·SE (where $\mathrm{t} = \mathrm{t}_{0.75,df}$ and $SE = SD/\sqrt{n}$ ). This yielded lower and upper bounds around the mean, with values below the lower bound classified as minimum activity, values within the bounds classified as average activity, and values above the upper bound classified as maximum activity within each timescale (figure 4). This classification provides a mean-centered measure of relative deviation in activity rates while accounting for sampling variability. All analyses were computed in RStudio v. 2023.6.0.421.

### S2 Theoretical Discussions

#### S2.1 Mean Relative Speed (MRS) Discussion

We begin by examining the phase-plane geometry of 4 scenarios under the **Mean Relative Speed** (MRS) assumption, comparing periodically forced activity patterns that are synchronous and asynchronous under slow and fast mean activity rates (a=0.25 and 0.7 respectively). As the periodic attack rate ($\alpha$) varies in response to these activity patterns, the interior equilibrium shifts across the phase-plane between the minimum ($\alpha_{\mathrm{min}}$) and maximum ($\alpha_{\mathrm{max}}$) deterministic attractors (i.e., stable equilibrium points and/or stable limit cycles, figure S2.1B), and the forced C-R dynamics track this time-varying interior equilibrium. We first note that the interior equilibria established by the mean attack rate ($\alpha_{\mathrm{mean}}$) are stable when the mean activity rate is slow (a=0.25;

figure S2.1A, C-R isoclines 1 and 2), and unstable when the mean activity rate is fast (a=0.7; figure S2.1A, C-R isoclines 3 and 4). On top of this, the mean attack rate increases with asynchronous activity patterns, resulting in the underlying C-R dynamics to become more excitable. Second, under asynchronous activity patterns, the periodic attack rate's amplitude is reduced while its frequency at which it oscillates at is doubled (figure 5Aii). This constrains the interior equilibrium to move over a smaller range within the phase-plane but over a shorter period of time, such that the forced C-R dynamics effectively average over the fluctuating interior equilibrium, resulting in trajectories that remain close to the mean equilibrium when activity patterns are asynchronous compared to when they are synchronous (i.e., the $\alpha_{\text{mean}}$ C-R isocline; see figure S2.1B2 and S2.1B1, respectively).

When the mean activity rate is slow (a=0.25), the equilibria corresponding to the mean attack rate are stable under both synchronous and asynchronous activity patterns (figure S2.1A – 1 and 2 respectively). Transitioning from synchronous to asynchronous activity patterns increases the consumer's mean attack rate ($\alpha_{\text{mean}}$) from 0.35 to 0.43 (see figure S2.1B1 and S2.1B2 insets respectively), which stabilizes the dynamics by shifting the consumer's mean density further away from near zero densities. In addition, the doubled oscillation frequency and the reduced range over which the interior equilibrium moves between the minimum ($\alpha_{\text{min}}$) and maximum ($\alpha_{\text{max}}$) deterministic attractors decreases the consumer's SD ($\alpha_{\text{SD}}$) from 0.25 to 0.05 (see figure S2.1B1 and S2.1B2 insets respectively), further enhancing stability. As a result, because the interior equilibrium now moves across a smaller range over a shorter period of time under asynchronous activity patterns (figure S2.1B2), the forced C-R dynamics are more strongly drawn towards the $\alpha_{\text{mean}}$ equilibrium. Under synchronous activity patterns, the dynamics are more influenced by the moving attractor and are pulled towards the minimum deterministic attractor due to asymmetric equilibrium tracking over this range (figure S2.1B1). Overall, under slow mean activity rates, increases in the mean attack rate, along with higher frequency but reduced-amplitude attack rate variation (and therefore equilibrium motion), collectively act together to stabilize the forced C-R dynamics under asynchronous activity patterns (figure 5Aiii).

When the mean activity rate is fast (a=0.7), the equilibria corresponding to the mean attack rate are unstable under both synchronous and asynchronous activity patterns (figure S2.1A – 3 and 4 respectively). As a result, the deterministic C-R dynamics exhibit oscillatory behaviour, with limit cycles at $\alpha_{mean}$ and $\alpha_{max}$ as the underlying attractors. Under periodic forcing, transitioning from synchronous to asynchronous activity patterns again increases the mean attack rate ($\alpha_{mean}$) from 0.99 to 1.20 (see figure S2.1B3 and S2.1B4 insets respectively), further elevating the consumer's attack rate on the resource and thereby acting as a destabilizing force on the system. Although asynchronous activity increases the periodic attack rate's oscillation frequency and reduces its variability ($\alpha_{SD}$ decreases from 0.7 to 0.14; see figure S2.1B3 and S2.1B4 insets respectively), which constrains the range in which the interior equilibrium moves and pulls the dynamics more strongly toward the $\alpha_{mean}$ limit cycle, this limit cycle has a larger amplitude under asynchronous conditions, resulting in greater population variability. In contrast, when activity patterns are synchronous, the interior equilibrium fluctuates between a stable equilibrium point ($\alpha_{min}$) and an unstable node ($\alpha_{max}$, now a stable limit cycle) beyond the Hopf bifurcation. Since the $\alpha_{mean}$ attractor is a limit cycle, the interior equilibrium spends more time in the oscillatory region of parameter space, thus inducing additional oscillatory behaviour within the forced C-R dynamics. However, the dynamics are additionally being pulled towards the stable minimum attractor due to asymmetric equilibrium tracking over this range (figure S2.1B3). Therefore, despite our forcing acting in qualitatively similar ways to the slow activity case, we see that asynchrony has an opposite effect on stability (consumer CV) when the mean activity rate is fast – that is, now asynchrony destabilizes the system. In this case, lower amplitude but higher frequency attack rate variation again causes the dynamics to become "averaged" around the mean deterministic attractor, however this attractor is now a larger-amplitude limit cycle (figure 5Aiv).

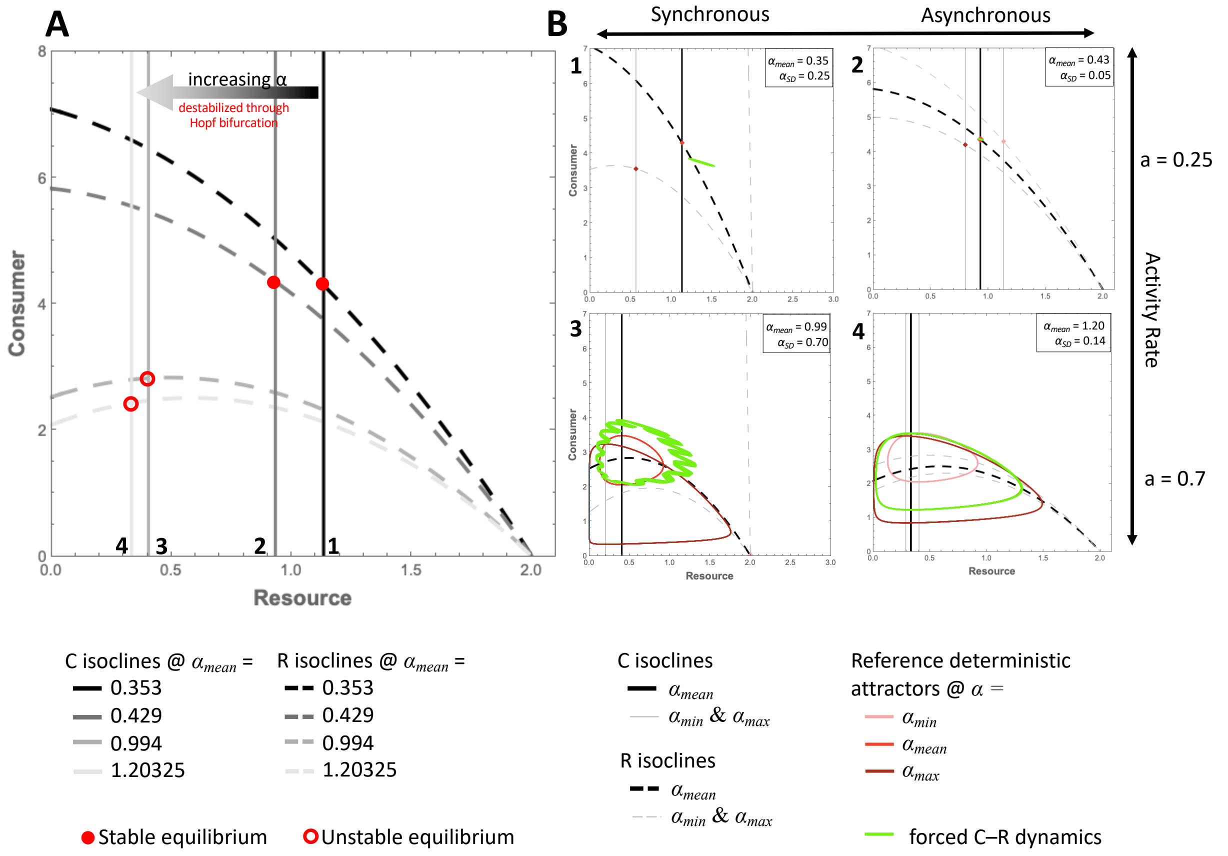


**Figure S2.1:** Graphical representation of changing the attack rate ($\alpha$) in a Rosenzweig-MacArthur consumer (C)-resource (R) model when activity patterns scale with the mean relative speed (MRS) assumption. A) The isoline geometry and equilibrium structure changes with increasing the mean attack rate $\alpha_{mean}$; and B) The underlying isocline geometry around each mean attack rate (in A) as it fluctuates between its respective minimum and maximum isoclines, and the behaviour of the forced C-R dynamics (from 900 to 1000 time-units) as activity patterns transition from synchrony (left panel) to asynchrony (right panel) under slow (top panel) and fast (bottom panel) mean activity rates.

S2.2 Mean Relative Speed with Refugia (MRS + Refugia) Discussion

We now examine the phase-plane geometry of 4 scenarios under the **Mean Relative Speed + Refugia** (MRS + Refugia) assumption when periodically forced activity patterns are synchronous and asynchronous under slow and fast mean activity rates (a=0.25 and 0.7 respectively). Again, the forced C-R dynamics attempt to track the interior equilibrium as it fluctuates between the minimum and maximum deterministic attractors in response to the fluctuating attack rate (i.e., via the fluctuating activity patterns; figure S2.2B). There are two notable differences for the MRS + Refugia assumption, compared to our previous MRS assumption, in terms of how they affect the attack rate variation. First, the refugia assumption gives a larger influence to the resource activity, particularly at low activity, making asynchronous activity patterns decrease the mean attack rates. Second, asynchrony no longer changes the frequency of attack rate variation. Here, under the MRS + Refugia assumption, asynchronous activity patterns are stabilizing under both slow and fast activity rates (figure 5Biii, 5Biv), with the stabilizing effect more pronounced at fast activity rates.

When activity rates are slow (a=0.25), the forced C-R dynamics become more stable as activity patterns transition from completely synchronous to completely asynchronous (figure 5Biii). Although this transition reduces the mean attack rate ($\alpha_{\text{mean}}$) from 0.33 to 0.26 (see figure S2.2B1 and S2.2B2 insets respectively), thereby lowering the already small consumer's mean density (a destabilizing force; figure S3.7a), it also reduces variability in the attack rate ($\alpha_{\text{SD}}$ decreases from 0.26 to 0.18; see figure S2.2B1 and S2.2B2 insets respectively; figure S3.7b) and consumer dynamics, such that this variance-driven stabilization counteracts the mean-driven destabilization. In both scenarios, the minimum and maximum deterministic attractors ($\alpha_{\text{min}}$ and $\alpha_{\text{max}}$) in which the interior equilibrium fluctuates between are stable. Here, the forced C-R dynamics primarily oscillate in the resource (R) direction while they respond to the fluctuating interior equilibrium. Under asynchronous activity patterns, the dynamics occupy a smaller region of the phase-plane due to the reduced variability in the attack rate. In both synchronous and asynchronous cases, the dynamics are also influenced by the moving interior equilibrium and are drawn more strongly towards the minimum deterministic attractor due to asymmetric equilibrium tracking over this range (figure S2.2B1 and S2.2B2).

Under fast mean activity rates (a=0.7), reductions in both the mean attack rate ($\alpha_{\text{mean}}$ from 0.94 to 0.73), and its variability ($\alpha_{\text{SD}}$ from 0.73 to 0.51; see S2.2B3 and S2.2B4 insets respectively) act together to stabilize the forced C-R dynamics as activity patterns transition from synchrony to asynchrony. The lower mean attack rate stabilizes the mean deterministic attractor ($\alpha_{\text{mean}}$) as it is shifted past the Hopf bifurcation (figure S2.2A – 3 to 4). Under synchronous activity patterns (figure S2.2B3), the interior equilibrium fluctuates between a stable equilibrium point ($\alpha_{\text{min}}$) and an unstable node ($\alpha_{\text{max}}$, now a limit cycle), crossing the Hopf bifurcation and traversing a wide region of the phase-plane due to the large attack rate variability. Moreover, because the $\alpha_{\text{mean}}$ attractor is also a limit cycle, the interior equilibrium spends a disproportionate amount of time in the oscillatory region of parameter space, thereby amplifying oscillations in the forced C-R dynamics. In contrast, under asynchronous activity patterns (figure S2.2B4), the interior equilibrium again fluctuates between a stable equilibrium point ($\alpha_{\text{min}}$) and a limit cycle ($\alpha_{\text{max}}$), but the $\alpha_{\text{mean}}$ attractor is now a stable point. This forces the interior equilibrium to spend more time in the stable equilibrium region of parameter space, and, together with the reduced mean and variability of the attack rate, the forced C-R dynamics are more constrained, remaining slightly above the $\alpha_{\text{mean}}$ deterministic attractor and varying primarily along the resource (R) axis. Combined with the reduction of the attack rate’s mean and variance, asynchronous activity patterns aid in stabilizing forced C-R dynamics when activity rates are fast under the MRS + Refugia assumption.

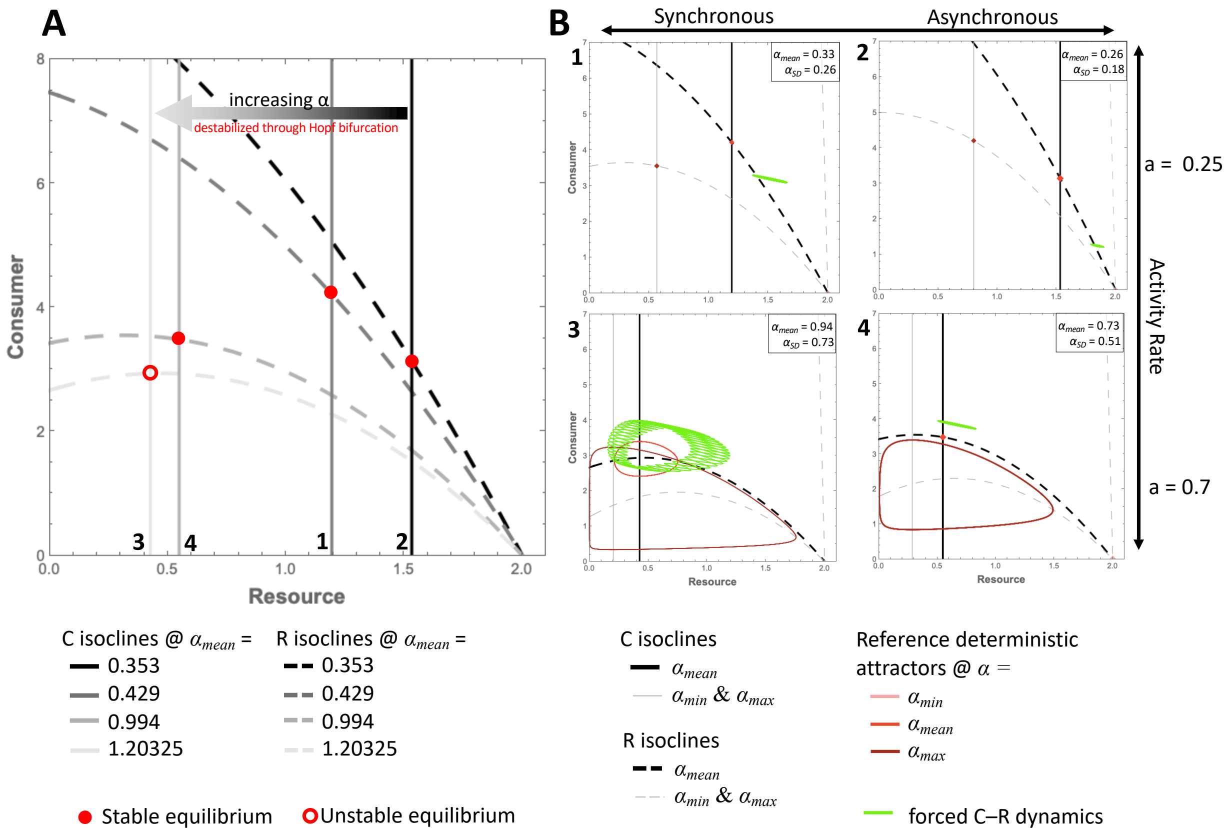


**Figure S2.2:** Graphical representation of changing the attack rate ($\alpha$) in a Rosenzweig-MacArthur consumer (C)-resource (R) model when activity patterns scale with the mean relative speed + refugia assumption (MRS + Refugia). A) The isoline geometry and equilibrium structure changes with increasing the mean attack rate $\alpha_{mean}$; and B) The underlying isocline geometry around each mean attack rate (in A) as it fluctuates between its respective minimum and maximum isoclines, and the behaviour of the forced C-R dynamics (from 900 to 1000 time-units) as activity patterns transition from synchrony (left panel) to asynchrony (right panel) under slow (top panel) and fast (bottom panel) mean activity rates.

**S3 Additional Supplementary Section (Figures)**

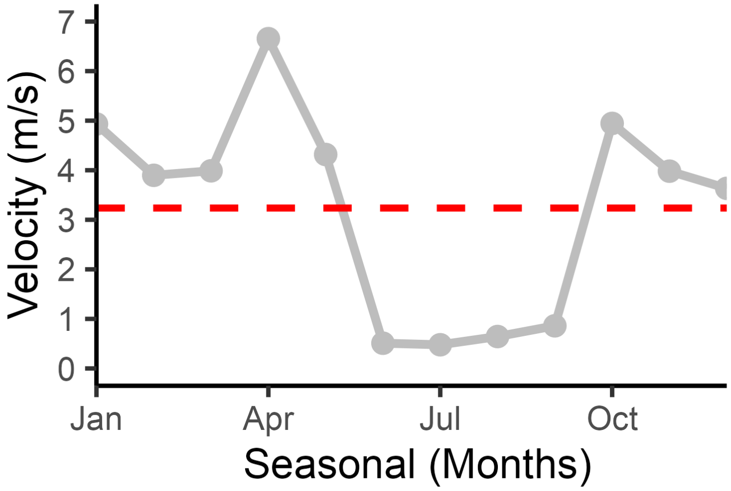


**Figure S3.1:** Full profile of great cormorant's (*Phalacrocorax carbo*) seasonal activity patterns. Larger mean of 3.237 m/s is due to the species migration periods spanning from October to April. Original sources for all data can be found in table S2, and each species' tagging duration can be found in table S3.

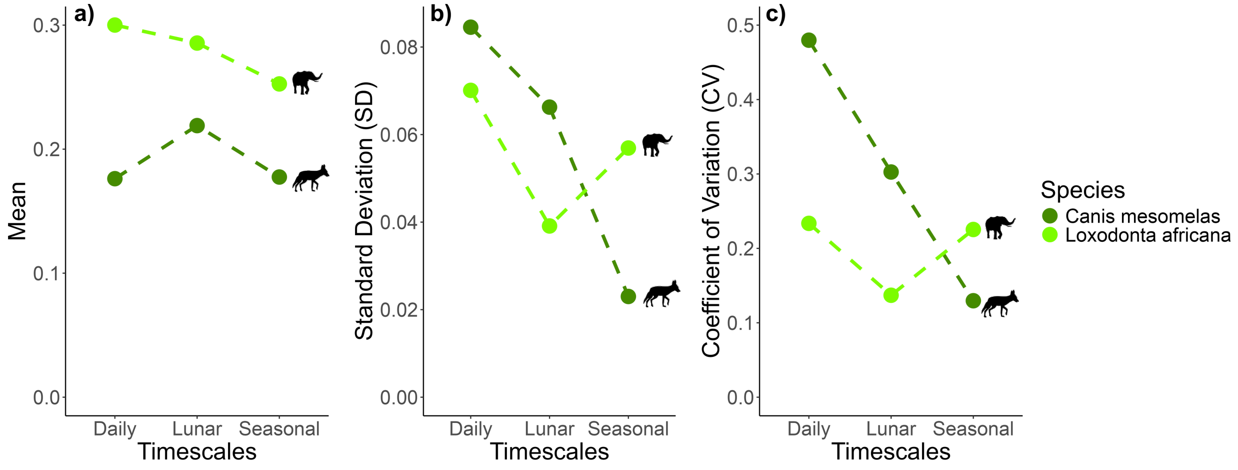


**Figure S3.2:** Close up on the two terrestrial species differing in their activity a) mean, b) standard deviation, and c) coefficient of variation across the daily, lunar, and seasonal timescales.

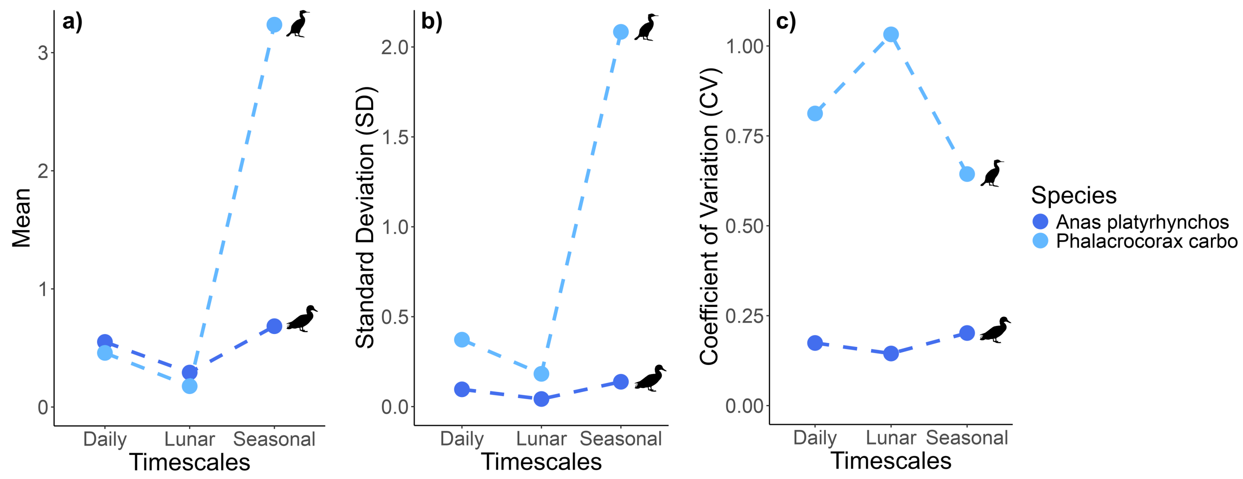


**Figure S3.3:** Close up on the two aerial species differing in their activity a) mean, b) standard deviation, and c) coefficient of variation across the daily, lunar, and seasonal timescales.

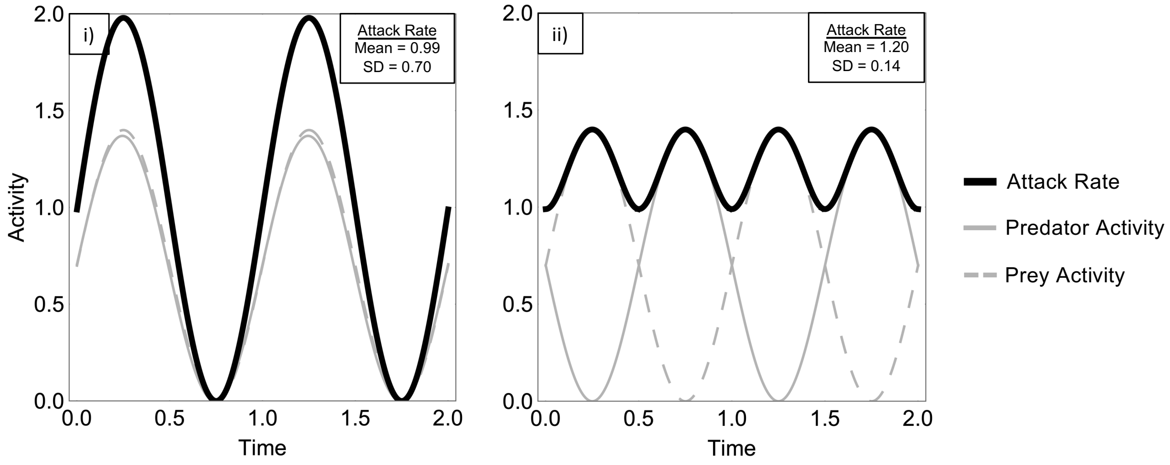


**Figure S3.4:** Relative maximum attack rate scales with the activity patterns of the predator and prey when the mean activity rate is fast (a=0.7) and the activity patterns are synchronous (i: *Ps* = 0) and asynchronous (ii: *Ps* = 1). Activity variation is the scaler percentage (*per*) of the predator and prey's activity. Parametric values: $r = 2.5$, $k = 2.0$, $m = 0.2$, $h = 1$, $e = 0.7$, $p = 1$.

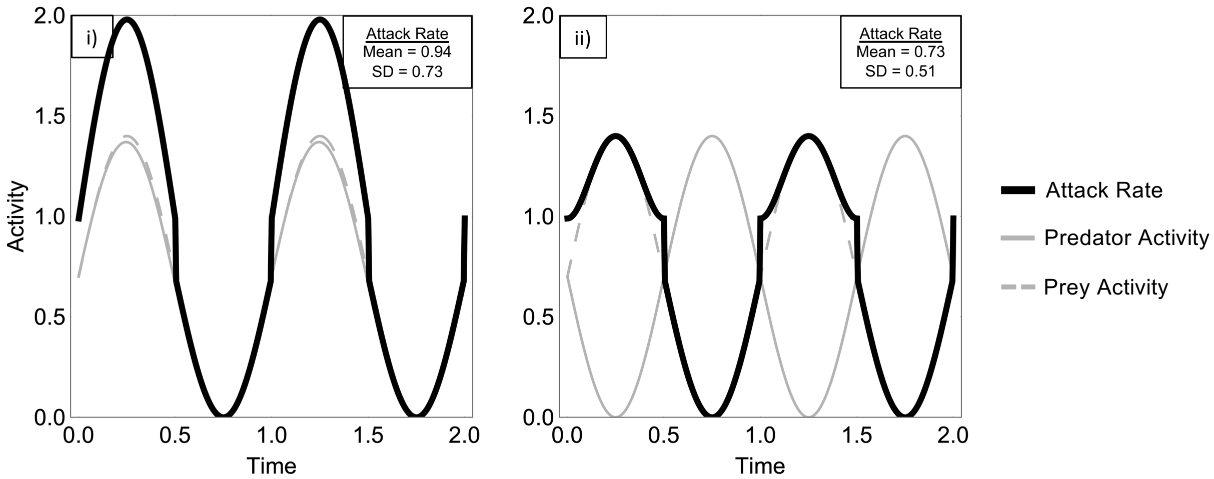


**Figure S3.5:** Relative maximum attack rate with refugia – scales with the activity patterns of the predator and prey when the mean activity rate is fast (a=0.7) and the activity patterns are synchronous (i: *Ps* = 0) and asynchronous (ii: *Ps* = 1). Activity variation is the scaler percentage (*per*) of the predator and prey's activity. Parametric values: $r = 2.5$, $k = 2.0$, $m = 0.2$, $h = 1$, $e = 0.7$, $p = 1$.

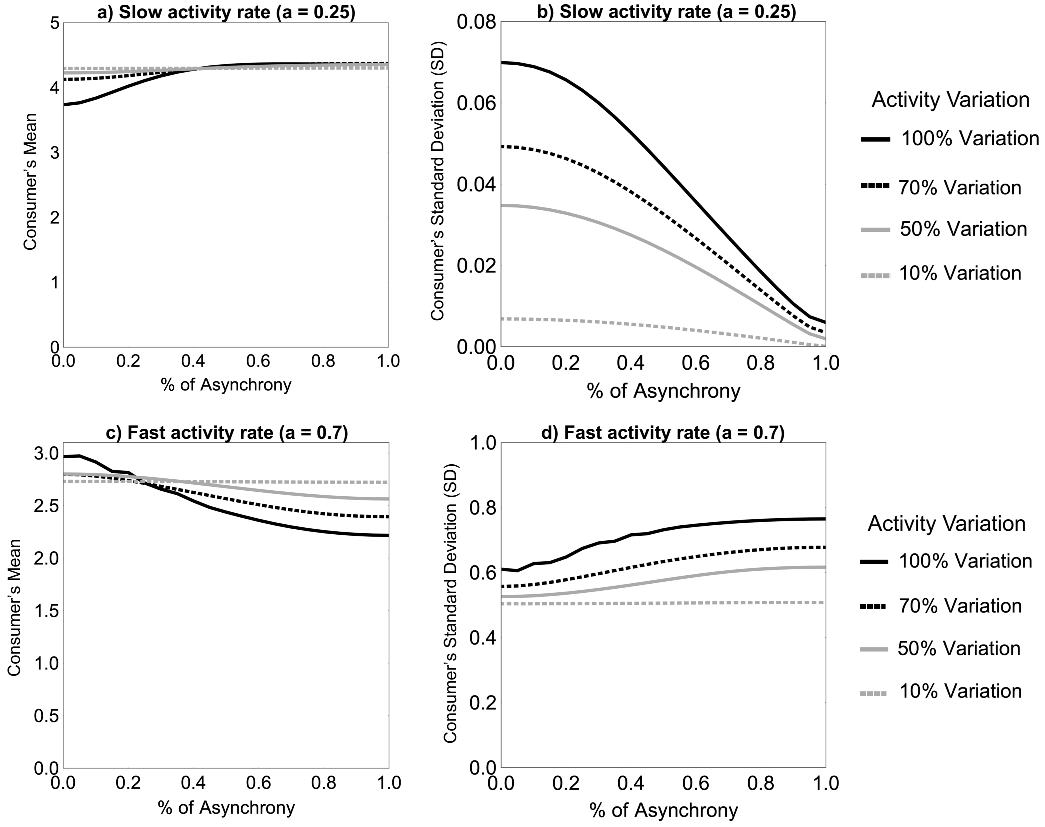


**Figure S3.6:** Predator's **a)** mean and **b)** standard deviation when the attack rate scales with a relatively slower mean activity rate (a=0.25). Predator's **c)** mean and **d)** standard deviation when the attack rate scales with a relatively faster mean activity rate (a=0.7).

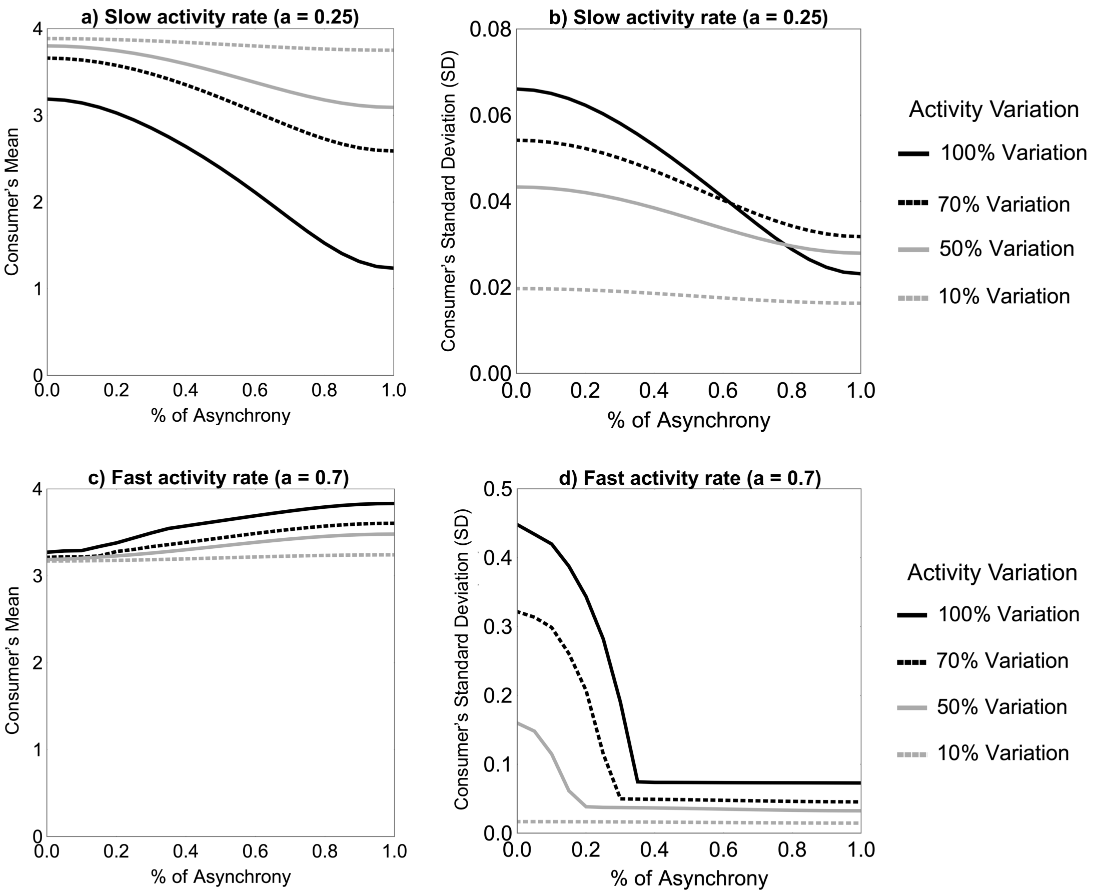


**Figure S3.7:** Predator's **a)** mean and **b)** standard deviation when the attack rate scales with a relatively slower mean activity rate with refugia (a=0.25). Predator's **c)** mean and **d)** standard deviation when the attack rate scales with a relatively faster mean activity rate with refugia (a=0.7).

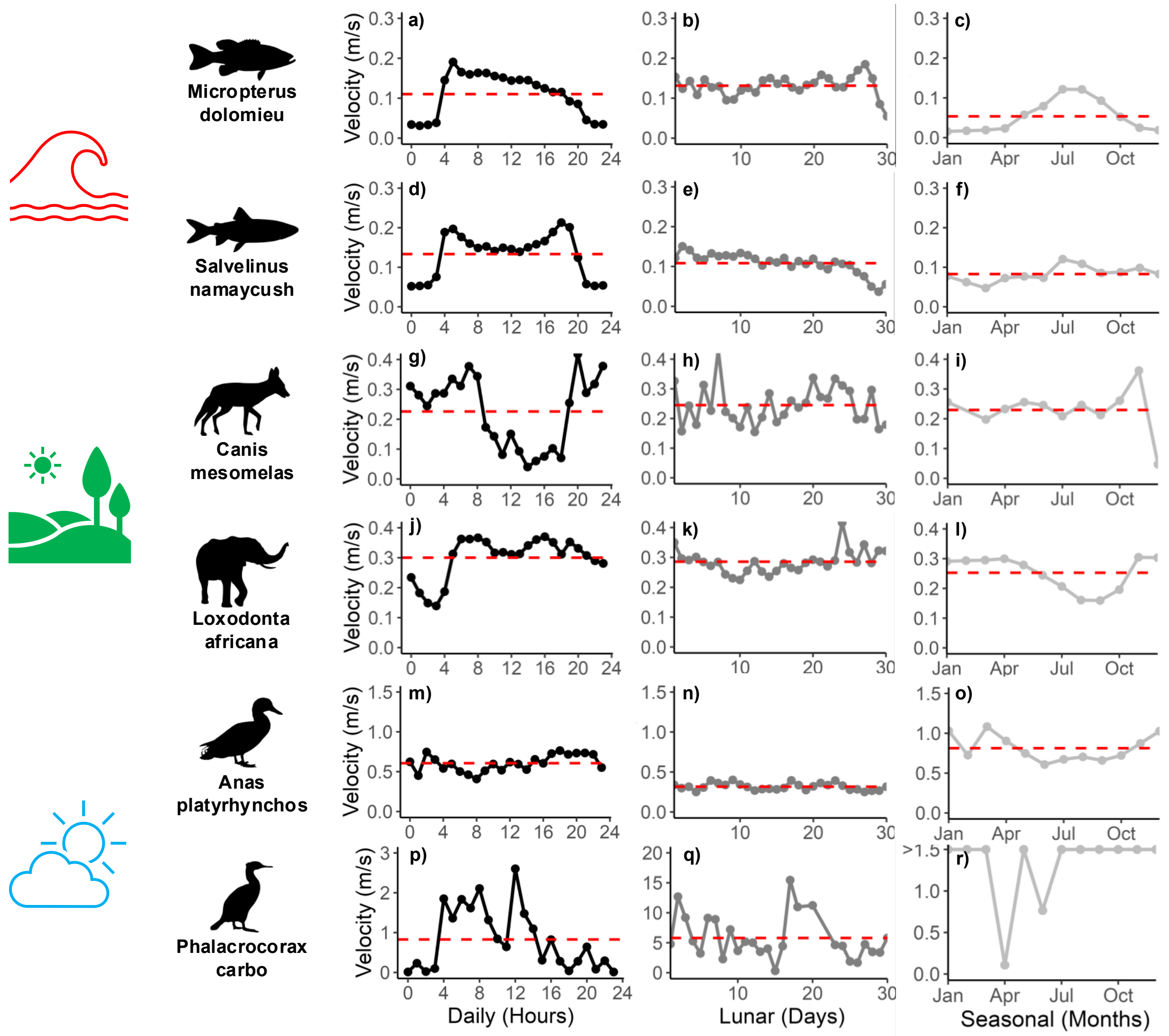


**Figure S3.8:** Activity patterns of various species across the daily (black lines), lunar (grey lines), and seasonal (light-grey lines) timescales with their corresponding activity mean (red dashed line) at tracking resolutions ≤ 28min intervals. Small mouth bass (*Micropterua dolomieu*; a-c) and lake trout (*Salvelinus namaycush*: d-e) are both aquatic cohabitants from Algonquin Park, Ontario Canada. Both terrestrial species, the black-backed jackal (*Canis mesomelas*: g-i) and the African elephant (*Loxodonta africana*: j-l), reside in Etosha National Park, Namibia, Africa, and the mallard duck (*Anas platyrhynchos*) and the great cormorant (*Phalacrocorax carbo*) are both aerial birds found near Lake Constance, Baden-Württemberg, Germany. Original sources for all data can be found in table S2, and each species' tagging duration can be found in table S4.

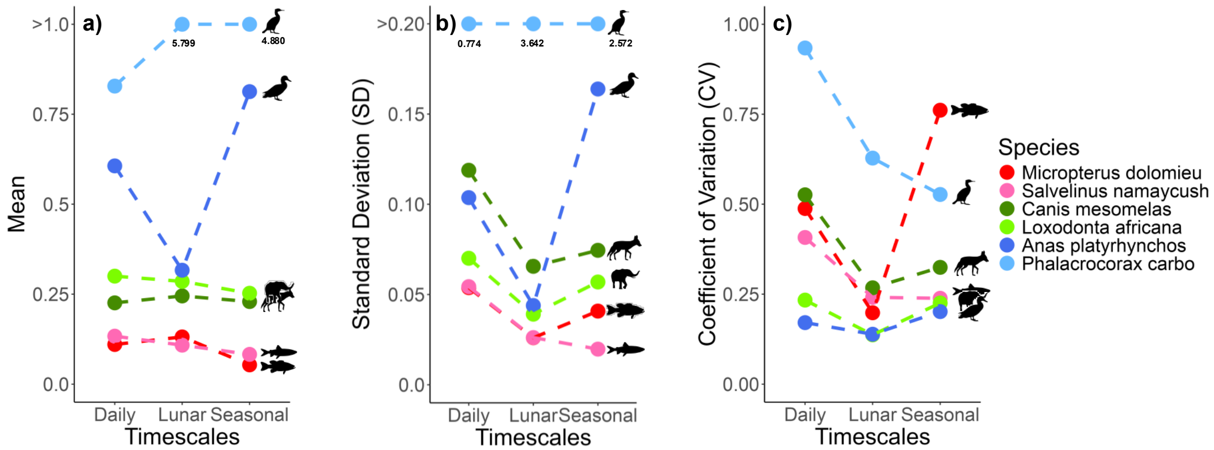


**Figure S3.9:** Relatively faster and slower activity patterns (measured by **(a)** mean) and variability of the activity patterns (measured by **(b)** standard deviation (SD) and **(c)** coefficient of variation (CV)) of various species across the daily, lunar, and seasonal timescales. All species are at a tracking resolution ≤ 28min interval. Original sources for all data can be found in table S2.

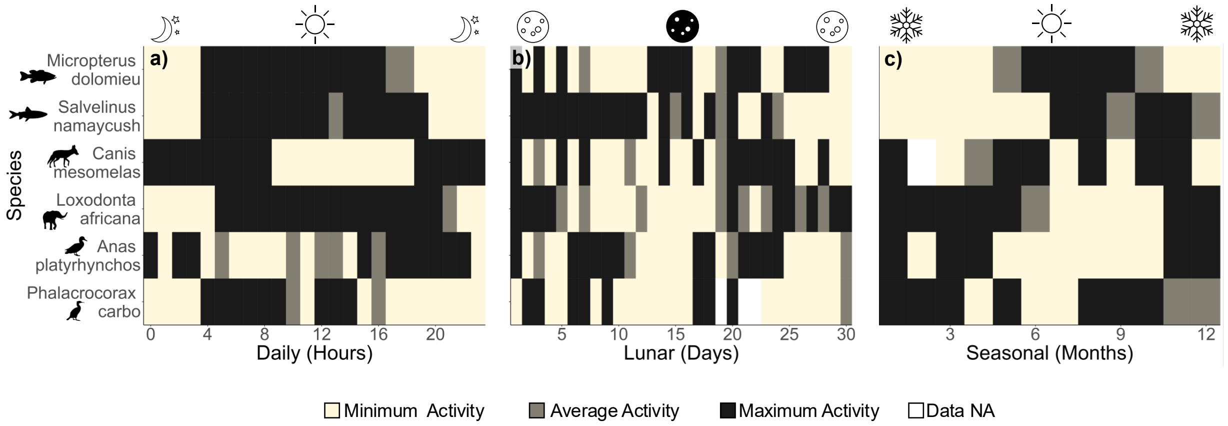


**Figure S3.10:** Times of minimum to maximum activity rates of various species across the (a) daily, (b) lunar, and (c) seasonal timescales, measured at 50% CI where the minimum activity is < 25% CI, average activity is from 25-75% CI, and maximum activity is > 75% CI. Species show both synchrony and asynchrony of activity rates at all scales. All species are at a tracking resolution ≤ 28min interval. Original sources for all data can be found in table S2.

**Tables**

**Table S1:** Predator-prey activity patterns

| Time Scale | Pred-prey timing | Variation | Predator | Peak predator activity | Prey | Peak prey activity | Location | Citation |
|---|---|---|---|---|---|---|---|---|
| Daily | Synch | Both variable | Coyotes | Crepuscular | Cottontail & Jackrabbit | Nocturnal & Crepuscular | Mapimí Biosphere Reserve (26° 40'N, 103°40'W), Mexico | Arias-Del Razo et al. (2011) |
| Daily | Synch | Both variable | Brown shrimp | Nocturnal | Juvenile plaice | Nocturnal & Crepuscular | Dunstaffnage Bay (56°30'N, 5°20'W), Scotland | Burrows (1994) |

| | | | | | | | | |
|---|---|---|---|---|---|---|---|---|
| Daily | Synch | All variable | Fox | Nocturnal | Hare & Rabbit | Nocturnal | Northern Ireland | Caravaggi et al. (2018) |
| Daily | Synch | All variable | Pine marten | Nocturnal | Wood mouse | Nocturnal | Northern Ireland | Caravaggi et al. (2018) |
| Daily | Asynch | All variable | Pine marten | Nocturnal | Squirrel | Diurnal | Northern Ireland | Caravaggi et al. (2018) |
| Daily | Asynch | Both variable | (Rare) Tasmanian Devil | Diurnal | Tasmanian pademelon, Bennett's wallaby, Common wombat | Crepuscular | Maria Island, Tasmania | Cunningham et al. (2019) |
| Daily | Synch | Both variable | (Abundant) Tasmanian Devil | Crepuscular | Tasmanian pademelon, Bennett's wallaby, Common wombat | Crepuscular | Mainland Tasmania | Cunningham et al. (2019) |

| | | | | | | | | |
|---|---|---|---|---|---|---|---|---|
| Daily | Asynch | Both variable | Hyenas | Nocturnal | Wildebeest | Crepuscular | Greater Serengeti-Mara, East Africa | de Jonge et al. (2022) |
| Daily | Asynch | Variable predator, constant prey | Hyenas | Nocturnal | Zebra | Crepuscular & Cathemeral | Greater Serengeti-Mara, East Africa | de Jonge et al. (2022) |
| Daily | Synch | Both variable | Tige | Nocturnal & Crepuscular | Wild boar | Nocturnal & Crepuscular | Hunchun Nature Reserve, Eastern Jilin Province, China | Dou et al. (2019) |
| Daily | Asynch | Both variable | Tiger | Nocturnal & Crepuscular | Roe and sika deer | Diurnal | Hunchun Nature Reserve, Eastern Jilin Province, China | Dou et al. (2019) |

| | | | | | | | | |
|---|---|---|---|---|---|---|---|---|
| Daily | Partially Synch | Both variable | Coyotes | Crepuscular | Does | Diurnal & crepuscular | Eastern Roan Mountain Highlands, North Carolina and Carter County, Tennessee, USA (36°6.34′N, 82°5.96′E) | Higdon et al. (2019) |
| Daily | Synch | Both variable | Coyotes | Crepuscular | Bucks | Crepuscular | Eastern Roan Mountain Highlands, North Carolina and Carter County, Tennessee, USA (36°6.34′N, 82°5.96′E) | Higdon et al. (2019) |

| | | | | | | | | |
|---|---|---|---|---|---|---|---|---|
| Daily | Asynch | Both variable | Coyotes | Crepuscular | Nursery Groups | Diurnal | Eastern Roan Mountain Highlands, North Carolina and Carter County, Tennessee, USA (36°6.34′N, 82°5.96′E) | Higdon et al. (2019) |
| Daily | Asynch | Both variable | Coyote, bobcat, bear, wolf | Nocturnal | Deer | Crepuscular | Upper Peninsula, Michigan, USA (46.54 N, 88.77 W) | Kautz et al. (2022) |
| Daily | Partially Synch | All variable | Hawks, kestrel, buzzard | Crepuscular (dawn) & diurnal | Sparrow, finches, robin, tits etc. | Crepuscular (dawn) | Great Britain/Ireland & North America | Lang et al. (2019) |

| | | | | | | | | |
|---|---|---|---|---|---|---|---|---|
| Daily | Synch | Both variable | Polecat | Nocturnal (spring & summer), crepuscular (fall & winter) | Amphibians: agile frog & common toad | Nocturnal | River Loire, Western France | Lode (1995) |
| Daily | Synch | Both variable | Polecat | Nocturnal (spring & summer), crepuscular (fall & winter) | Rodents: common vole, bank vole & brown rat | Crepuscular | River Loire, Western France | Lode (1995) |
| Daily | Synch (winter, summer), Asynch | Both variable | Wildcat | Nocturnal (winter, spring, autumn), | Rabbits | Cathemeral (winter), Crepuscular (spring, | Sierra Harana, Betic Mountains, Granada province, Spain | Martín-Díaz et al. (2018) |

| | | | | | | | | |
|---|---|---|---|---|---|---|---|---|
| | (Spring, Autumn) | | | Crepuscular (summer) | | summer), Diurnal (summer) | | |
| Daily | Asynch | Both variable | Great barracuda | Diurnal & Crepuscular | Schoolmaster snapper | Nocturnal | Guanica Biosphere, Puerto Rico | Rooker et al. 2018 |
| Daily | Asynch | Both variable | Great barracuda | Diurnal & Crepuscular | White grunt | Crepuscular | Guanica Biosphere, Puerto Rico | Rooker et al. 2018 |
| Daily | Synch | Both variable | Coopers hawks | Diurnal & Crepuscular | European starlings, mourning doves, and rock pigeons | Crepuscular | Western Indiana, USA | Roth and Lima (2007) |
| Daily | Partially asynch | Both variable | Sharp shinned hawks | Diurnal (sunset) | European starlings, | Crepuscular | Western Indiana, USA | Roth and Lima (2007) |

| | | | | | | | | |
|---|---|---|---|---|---|---|---|---|
| | | | | | mourning doves, and rock pigeons | | | |
| Daily | Asynch | Variable prey, constant predator | Lynx | Cathemeral | Snowshoe hare | Nocturnal | Southern Yukon, Canada (61°N, 138°W) | Shiratsuru et al. (2023) |
| Daily | Asynch | All variable | Lion | Nocturnal & Crepuscular | Warthog, kudu & elephant | Diurnal | Addo Elephant National Park, South Africa | Tambling et al. (2015) |
| Daily | Synch | Both variable | Lion | Nocturnal & Crepuscular | Buffalo | Diurnal & crepuscular | Addo Elephant National Park, South Africa | Tambling et al. (2015) |
| Daily | Asynch | All variable | Hyaena | Nocturnal | Warthog, kudu & elephent | Diurnal | Addo Elephant National Park, South Africa | Tambling et al. (2015) |

| | | | | | | | | |
|---|---|---|---|---|---|---|---|---|
| Daily | Asynch | Both variable | Hyaena | Nocturnal | Buffalo | Diurnal & crepuscular | Addo Elephant National Park, South Africa | Tambling et al. (2015) |
| Daily | Synch | Both variable | Saharan sand viper | Nocturnal (cathemeral) | Wedge snouted skink, Lichtenstein's short-fingered gecko, Anderson's short-fingered gecko | Nocturnal (cathemeral) | Desert sand dunes, Western Negev, Israel | Tesler et al. (2023) |
| Daily | Asynch | Both variable | Saharan sand viper | Nocturnal (cathemeral) | Egyptian fringe-fingered lizard, sandfish | Diurnal | Desert sand dunes, Western Negev, Israel | Tesler et al. (2023) |
| Daily | Synch | Both variable | Wolves | Peak at dusk | Moose | Crepuscular | Northern Ontario, Canada | Vander Vennen et al. (2016) |

| | | | | | | | | |
|---|---|---|---|---|---|---|---|---|
| Lunar | Synch | All variable | Parnell's mustached bat, Greater sac-winged bat, Lesser sac-winged bat | Full Moon | Beetles | Full moon | Reserva Florestal Adolpho Ducke (2°58′S, 59°55′W), Brazil. | Appel et al. (2017) |
| Lunar | Synch | Both variable | Short-eared owl | New moon | Deer mice | New moon | Laboratory | Clarke (1983) |
| Lunar | Asynch | variable | Jaguar | First quarter/full moon | Sea turtles (on land activity) | Last quarter/ new moon | Corcovado National Park, Costa Rica | Carrillo et al. (2009) |
| Lunar | Synch | All variable | Ocelot | 3rd moon | White-eared opposum, Grey four-eyes opossum, Tapeti | 3rd moon | Estancia Guaycolec ranch, South America | Huck et al. (2017) |

| | | | | | | | | |
|---|---|---|---|---|---|---|---|---|
| | | | | | | | (58º11W, 25º58S) | |
| Lunar | Asynch | Both variable | Ocelot | 3rd moon | Nine-banded armadillo | New & full moon | Estancia Guaycolec ranch, South America (58º11W, 25º58S) | Huck et al. (2017) |
| **These below have multiple timescales in one paper** | | | | | | | | |
| Daily | Synch | Variable (predator & prey) | Wolf | Crepuscular in summer, diurnal in winter | Moose | Crepuscular in summer, diurnal in winter | Voyageurs National Park, Northern Minnesota, USA | Ditmer et al. (2018) |
| Seasonal | Synch | Constant | Wolf | Active in all studied seasons | Moose | Active in all studied seasons | | |

| | | | | | | | | |
|---|---|---|---|---|---|---|---|---|
| Daily | Asynch | Variable (predator & prey) | Snow leopard | Nocturnal (sunrise in summer), Nocturnal (sunset in winter) | Ibex | Diurnal & Crepuscular | | |
| Lunar | Synch | Constant (slightly negatively associated with moonlight in predator, constant in prey) | Snow leopard | | Ibex | | Tost Mountains (43°N, 100°E), Gobi Desert, Mongolia | Johansson et al 2022 |

| | | | | | | | | |
|---|---|---|---|---|---|---|---|---|
| Seasonal | Synch | Constant (predator & prey) | Snow leopard | Consistent all year | Ibex | Active all year, slightly higher in summer | | |
| Daily | Synch | Variable predator, constant prey | Yellow legged hornets | Diurnal (all day) | European honeybees | Diurnal (morning) | Experimental apiary (INRA, Villenave d'Ornon, France, 44º47N 0º34W) | Monceau et al. (2013) |
| Seasonal | Asynch | Variable | Yellow legged hornets | Early autumn | European honeybees | Early summer | | |
| Daily | Synch | Both variable | Lynx | Crepuscular | Rabbits | Crepuscular | Doñana National Park, Spain | Penteriani et al. (2013) |

| | | | | | | | | |
|---|---|---|---|---|---|---|---|---|
| Daily | Synch | Both variable | Red Fox (also a prey of lynx) | Nocturnal & Crepuscular | Rabbits | Crepuscular | | |
| Lunar | Synch | Prey more variable than predator | Red Fox (also a prey of lynx) | Slight increase at new moon | Rabbits | New moon | | |
| Daily | Synch | All variable | Ocelots | Nocturnal | Small mammal prey: armadillos, 3 opossums, spiny rat | Nocturnal & Crepuscular | | |
| Lunar | Asynch | Constant predator, variable prey | Ocelots | Did not respond to the moon | Armadillos & spiny rats | More active at new moon | Central Amazonia | Pratas-Santiago et al. (2016) |

| | | | | | | | | |
|---|---|---|---|---|---|---|---|---|
| Daily | Synch | All variable | Red fox, Stone marten, European badger, Common genet, & European wildcat | Nocturnal | Small mammals | Nocturnal | | |
| Seasonal | Synch | Variable (small mammal prey, red fox & stone marten), | Red fox, Stone marten, European badger, Common genet, & | Nocturnal | Small mammals | Nocturnal (activity duration increased & overlap with predators increased in | Montseny Natural Park, Catalan Pre-Coastal Range, North-East Iberian Peninsula | Vilella et al. (2020) |

| | | | |
|---|---|---|---|
| constant (badger, genet, wildcat) | European wildcat | | autumn & winter) |

**Table S2:** Original sources of where all animal tracking data and abiotic data can be found.

| Data | Location | Source | File (if applicable) |
|---|---|---|---|
| **Species** | | | |
| *Micropterus dolomieu* | Smoke Lake, Algonquin Provincial Park, ON, Canada (45.5172°N, -78.6816°W) | Unpublished | |
| *Salvelinus namaycush* | Smoke Lake, Algonquin Provincial Park, ON, Canada (45.5172°N, -78.6816°W) | Unpublished | |
| *Canis mesomelas* | Etosha National Park, Namibia, Africa (19°S,16°E) | Movebank | Blackbacked jackal, Etosha National Park, Namibia |
| *Loxodonta africana* | Etosha National Park, Namibia, Africa (19°S, 16°E) | Movebank | African elephant in Etosha National Park (data from Tsalyuk et al. 2018) |

| | | | |
|---|---|---|---|
| *Anas platyrhynchos* | Lake Constance, Baden-Württemberg, Germany (47.767ºN, 8.995ºE) | Movebank | Life Tracks Ducks Lake Constance |
| *Phalacrocorax carbo* | Lake Constance, Baden-Württemberg, Germany (47.694ºN, 9.005ºE) | Movebank | Great Cormorant Lake Constance MPIAB |
| **Abiotic Conditions** | | | |
| Temperature | Algonquin Park East Gate, Ontario (45.53°N, -78.27ºW) | Historical Climate Data, Environment and Climate Change Canada | |
| Snow Depth | Algonquin Park East Gate, Ontario (45.53°N, -78.27ºW) | Historical Climate Data, Environment and Climate Change Canada | |

| | | | |
|---|---|---|---|
| Solar Irradiance | Algonquin Park East Gate, Ontario (45.53°N, -78.27°W) | RStudio | getSunPosition {suncalc} |
| Moon Illumination | Algonquin Park East Gate, Ontario (45.53°N, -78.27°W) | RStudio | getMoonIllumination {suncalc} |

295

**Table S3:** The exact date ranges when all tracking data was used for the analyses of each species within each timescale. n represents the number of cycles (i.e., day, month (lunar), year) that were used in the manuscript analysis.

| **Species** | **Daily** | | **Lunar** | | **Seasonal** | |
|---|---|---|---|---|---|---|
| | **n** | **Dates** | **n** | **Dates** | **n** | **Dates** |
| *Micropterus dolomieu* | 31 | July 2021 | 1 | July 24th – August 23rd, 2021 | 1 | July 1st, 2021 – July 1st, 2022 |
| *Abiotic Conditions | 31 | July 2021 | 1 | July 24th – August 23rd, 2021 | 1 | July 1st, 2021 – July 1st, 2022 |
| *Salvelinus namaycush* | 31 | July 2021 | 1 | July 24th – August 23rd, 2021 | 1 | July 1st, 2021 – July 1st, 2022 |
| *Canis mesomelas* | 60 | April 2009<br>April 2010 | 2 | August 6th – September 4th, 2009<br>September 24th – October 23rd, 2010 | 2 | February 7th, 2009 – January 16th, 2011 |
| *Loxodonta africana* | 150 | | 5 | March 11th – April 9th, 2009<br>March 1st – 30th, 2010 | 5.5 | October 28th, 2008 – March 28th, 2014 |

| | | | | | | |
|---|---|---|---|---|---|---|
| | | April 2009,<br>2010, 2011,<br>2012, 2013 | | February 9th – March 20th, 2011<br>October 30th – November 28th, 2012<br>October 18th – November 18th, 2013 | | |
| *Anas platyrhynchos* | 150 | June 2017,<br>2018, 2019,<br>2020, 2021 | 4 | June 28th – July 27th, 2018<br>May 19th – June 17th, 2019<br>June 6th – July 5th, 2020<br>May 27th – June 25th, 2021 | 4.83 | January 8th, 2017 – November 10th, 2021 |
| *Phalacrocorax carbo* | 120 | June 2009,<br>2010, 2011<br>& 2012 | 1 | July 8th – August 6th, 2009 | 3 | June 17th, 2009 – July 4th, 2012 |

**Table S4:** The exact date ranges when all tracking data was used for the analyses of each species within each timescale where all species are at a resolution ≤ 28min between each fix interval. n represents the number of cycles (i.e., day, month (lunar), year) that were used in the analysis of supplementary figure S8, S9 and S10.

| Species | Daily n | Daily Dates | Lunar n | Lunar Dates | Seasonal n | Seasonal Dates |
|---|---|---|---|---|---|---|
| *Micropterus dolomieu* | 31 | July 2021 | 1 | July 24th – August 23rd, 2021 | 1 | July 1st,2021 – July 1st, 2022 |
| *Salvelinus namaycush* | 31 | July 2021 | 1 | July 24th – August 23rd, 2021 | 1 | July 1st,2021 – July 1st, 2022 |
| *Canis mesomelas* | 60 | April 2009<br>April 2010 | 2 | August 6th – September 4th, 2009<br>April 10th – May 9th, 2010 | 1.5 | March 8th, 2009 – October 8th, 2010 |
| *Loxodonta africana* | 150 | April 2009, 2010, 2011, 2012, 2013 | 5 | March 11th – April 9th, 2009<br>March 1st – 30th, 2010<br>February 9th – March 20th, 2011 | 5.5 | October 28th, 2008 – March 28th, 2014 |

| | | | | | | |
|---|---|---|---|---|---|---|
| | | | | October 30th – November 28th, 2012<br>October 18th – November 18th, 2013 | | |
| *Anas platyrhynchos* | 150 | June 2017, 2018, 2019, 2020, 2021 | 4 | June 28th – July 27th, 2018<br>May 19th – June 17th, 2019<br>June 6th – July 5th, 2020<br>May 27th – June 25th, 2021 | 4.83 | January 8th, 2017 – October 29th, 2021 |
| *Phalacrocorax carbo* | 120 | June 2009, 2010, 2011 & 2012 | 2 | July 7th – August 6th, 2009<br>August 6th – 29th, 2009 | 3 | June 17th, 2009 – July 4th, 2012 |

**Table S5:** Comparison between the total number of positions and number of individuals within the original datasets vs each filtered dataset when the resolution is ≤ 1hr and ≤ 28min between each fix point.

| **Species** | **Original Dataset (unfiltered)** | | **Main Analysis (Time elapsed: ≤ 1hr)** | | **Supplement (Time elapsed: ≤ 28min)** | | **Movebank Study Name** |
|---|---|---|---|---|---|---|---|
| | Total # of positions | Total # of individuals | # of positions | # of Individuals | # of positions | # of Individuals | |
| Micropterus dolomieu | 419150 | 18 | | | 419150 | 18 | |
| Salvelinus namaycush | 780218 | 26 | | | 780218 | 26 | |
| Canis mesomelas | 130688 | 22 | 88085 | 22 | 24907 | 12 | Black-backed jackal, Etosha National Park, Namibia |
| Loxodonta africana | 2930263 | 15 | 2160818 | 15 | 2160419 | 15 | African elephants in Etosha National Park |

| | | | | | | | |
|---|---|---|---|---|---|---|---|
| | | | | | | | (data from Tsalyuk et al. 2018) |
| Anas platyrhynchos | 1135291 | 72 | 957935 | 71 | 822550 | 70 | LifeTrack Ducks Lake Constance |
| Phalacrocorax carbo | 99836 | 11 | 6796 | 11 | 1930 | 10 | Great Cormorant Lake Constance |
